# Host-Botrytis co-transcriptomics reveals finely tuned interactions with closely related legumes

**DOI:** 10.64898/2026.01.20.700702

**Authors:** Anna Jo Muhich, Ritu Singh, Cloe Tom, Celine Caseys, Karishma Srinivas, Lucca Faieta, Brooke Grabbe, Daniel J. Kliebenstein

**Affiliations:** Department of Plant Sciences, University of California Davis, Davis CA 95616

**Keywords:** Comparative co-transcriptomics, legumes, necrotrophic fungal pathogenesis, network biology, transcriptional plasticity

## Abstract

Generalist pathogens infect diverse plant hosts, yet how these interactions differ across hosts is poorly understood. Here, we conduct a molecular analysis of a generalist pathogen interacting with closely related hosts. A co-transcriptomic framework is used to dissect host–pathogen interactions between the generalist necrotroph *Botrytis cinerea* and two closely related legume hosts, common bean (*Phaseolus vulgaris*) and cowpea (*Vigna unguiculata*). Using a diverse set of 72 Botrytis isolates, we quantified lesion development alongside host and pathogen gene expression. Although lesion formation was driven primarily by pathogen genetic variation, transcriptomic responses in both host and pathogen exhibited significant host x isolate interactions. This indicated that extensive, fine-scale transcriptional plasticity created similar disease outcomes. Botrytis genes showing host-specific expression were enriched for cell wall–modifying enzymes and some specialized metabolic genes, indicating greater host responsiveness of these core virulence mechanisms than previously appreciated. Co-expression network analysis in both host and pathogen further showed that in both organisms, gene membership for individual networks are restructured in response to genetic diversity. For example in Botrytis, we identify different sets of genes host-dependently co-expressing with a non-ribosomal peptide synthetase (NRPS) gene cluster, suggesting divergent functional deployment of the same virulence machinery across closely related hosts. Both legume species exhibited extensive isolate-dependent transcriptional reprogramming, with approximately two-thirds of expressed host genes responding to pathogen diversity. While conserved defense pathways such as jasmonate/ethylene signaling and phenylpropanoid metabolism were upregulated in both hosts, the specific genes in the networks differed markedly, highlighting lineage-specific rewiring of defense strategies. These results suggest that generalist pathogen success is underpinned by pervasive gene expression plasticity in both host and pathogen, allowing similar phenotypic outcomes to emerge from highly divergent molecular states.

**Summary:**

- Generalist pathogens infect diverse plant hosts, yet how these interactions differ across hosts is poorly understood. This study investigates how a generalist pathogen achieves successful infection across closely related hosts, and how these hosts respond.
- A co-transcriptomic approach was applied to interactions between 72 genetically diverse isolates of the fungal necrotroph *Botrytis cinerea* and two legume hosts, common bean and cowpea. Lesion development and host and pathogen gene expression were quantified.
- Lesion formation was primarily driven by pathogen genetic variation, yet both host and pathogen transcriptomes showed strong host x isolate interactions. Both host and pathogen balance conserved responses with finely tuned, host-specific mechanisms. Further, host-dependent transcriptional responses involve network modulation around a common core of genes in both host and pathogen.
- Generalist pathogen success is underpinned by pervasive gene expression plasticity in both host and pathogen, allowing similar phenotypic outcomes to emerge from highly divergent molecular states.

## Introduction

Plant species rely on a diverse range of defensive strategies to defend against pathogens. These defenses are frequently multi-functional allowing them to act against multiple pathogens or other biotic attackers. For example, chemical defenses like individual glucosinolates and monolignols, as well as more physical defenses like callose deposition simultaneously provide resistance to multiple bacterial and fungal pathogens, both generalists and specialists, as well as insect herbivores (Liu et al. 2021; Ninkuu et al. 2022; Wang et al. 2021). During plant speciation, diverging species will often occupy different environmental niches that likely differ in the challenging pathogen populations, creating pressure on each species to separately optimize their defense portfolio. This can include changes in their specific defense strategies and/or signaling pathways controlling these defenses. Across distantly related species, SA (salicylic acid) and JA (jasmonic acid) signaling generally functions as key response pathways to diverse biotic attackers. However, this does not mean that the entirety of the systems are conserved as the production of the signals, the connectivity between the pathways, and the distinct downstream outputs often vary dramatically (Wang et al. 2015; Monte 2023). While this defense system variation occurs across distantly related species, it is unclear how variable defense pathways and responses are between closely related plant species. To investigate this, we use a common generalist pathogen to stimulate and measure the defense responses in related plant species.

Evolutionary shifts in host plants raises a compelling question is how generalists successfully infect the broad range of evolving hosts they encounter. Generalists are predominantly thought to use mechanisms that function across a multitude of diverse hosts. These can include more promiscuous digestive enzymes (Li et al. 2004), effector proteins that induce necrosis (Barrett and Heil 2012), enzymes that neutralize reactive oxygen species (López-Cruz et al. 2017), or phytotoxic metabolites (Colmenares et al. 2002). However, to infect many hosts, a generalist must either respond to the diversity of specialized defenses and shifts in host defense signaling, or evade these defenses entirely. This suggests that successful generalists could rely on highly flexible or inducible counter-defense strategies that function across diverse host contexts. Somewhat counterintuitively, such flexibility implies generalists may deploy inducible specialized mechanisms that are expressed on specific hosts. There is some evidence for this in generalist pathogens, including specialized counter-defense efflux transporters (Stefanato et al. 2009). At large scale, other generalist pathogens have shown extensive transcriptional reprogramming when colonizing distinct hosts, including *Fusarium virguliforme* on maize and soybean (Baetsen-Young et al. 2020), *Sclerotinia sclerotium* on six eudicot families (Kusch et al. 2022), and *Botrytis cinerea* on ten eudicot species (Singh et al. 2025). However, given the diversity of eudicot hosts, it remains unclear how finely tuned a generalist pathogen’s virulence mechanisms are when comparing closely related host species infected by the same generalist pathogen.

To measure how host responses shift between closely related host plants and how a generalist pathogen compensates, we focused on the *Botrytis cinerea*-legume pathosystem. *Botrytis cinerea*, hereafter Botrytis, is a globally dispersed generalist fungal necrotroph with a host range spanning >1,400 land plant species, including vascular and nonvascular plants (Elad et al. 2016). Botrytis has extensive genome-wide sequence diversity that underlies complex quantitative disease outcomes across its many hosts (Rowe and Kliebenstein 2007; Fekete et al. 2012; Walker et al. 2015; Mercier et al. 2021; Caseys et al. 2021). To infect this diversity of hosts, Botrytis employs a wide array of virulence mechanisms, including phytotoxic metabolites, cell wall degrading enzymes, and cell death inducing proteins (Colmenares et al. 2002; Zhu et al. 2017), many of which exhibit substantial functional redundancy (Dalmais et al. 2011; Leisen et al. 2022). With a multitude of well described strategies, Botrytis provides an excellent system to understand how genetic variation within a generalist pathogen influences infection strategy, and how these strategies shift when challenging different plant hosts.

Among Botrytis’s many hosts are the Fabaceae or legume family in the rosids clade. Legumes are a widely grown crop and are of growing importance for sustainable agriculture (Voisin et al. 2014). Common bean (*Phaseolus vulgaris*) and cowpea (*Vigna unguiculata*) are two closely related agronomically important legumes, diverging ∼10 Mya (Moghaddam et al. 2021). As known Botrytis hosts, they provide a model for comparing host response specificity of closely related species when challenged with the same pathogen. Legumes represent the third largest plant family and display a massive diversity of potentially toxic specialized metabolites, including alkaloids, non-protein amino acids, cyanogens, peptides, phenolics, polyketides, and terpenoids (Wink 2013). In common bean and cowpea, both species likely use flavonoids and other phenolic compounds for defense, but comprehensive metabolomic studies for these species are minimal (Perez De Souza et al. 2019). Due to the large diversity of legume bioactive metabolites, both known and unknown, defensive strategies between these species likely diverge in the specialized metabolic pathways they utilize against the same pathogen.

Alongside growing recognition that gene expression plasticity contributes to the success of generalist pathogens, increasing evidence indicates that Botrytis also undergoes extensive transcriptional reprogramming in response to different hosts. When Botrytis gene expression was compared on wild-type *Arabidopsis thaliana* versus SA/JA signaling mutants, 45-74% of the detectable Botrytis transcriptome was differentially expressed across host genotypes (Zhang et al. 2019). Much of its expression variation was attributable to differences in Botrytis isolate and was largely controlled in trans, suggesting coordinated regulatory programs rather than gene-by-gene responses (Krishnan et al. 2023). Consistent with this, transcriptome profiling of 72 Botrytis isolates across 15 eudicot species identified sets of Botrytis genes with expression unique to individual host species (Singh et al. 2025). Together, these results indicate that Botrytis deploys distinct, host-dependent transcriptional programs during infection, raising the question of how hosts respond to and potentially shape these pathogen strategies. In some fungi including Botrytis, it is possible to simultaneously measure host and pathogen transcriptomes during infection, using a co-transcriptomic approach. This framework has been previously applied in Arabidopsis, revealing correlated host-pathogen gene co-expression networks and identifying novel mechanisms underlying the Arabidopsis-Botrytis interaction (Zhang et al. 2019). Co-transcriptomics therefore provides a powerful platform for dissecting the reciprocal strategies of host and pathogen during infection and can be extended to other agronomically important hosts of Botrytis.

In this study, we used the legume-Botrytis pathosystem to map the host-specific aspects shaping the interaction when comparing closely related hosts. This involved first measuring Botrytis virulence on different genotypes of legumes. Lesion sizes were quantified using 72 diverse *B. cinerea* isolates on 4 genotypes each of common bean and cowpea. For a molecular investigation, the co-transcriptomes of all 72 diverse Botrytis isolates and 1 genotype each of common bean and cowpea was measured during infection (Figure 1). This provided a map of how gene expression patterns in host and pathogen contribute to virulence, and how these patterns may change across genetically diverse individuals. This allowed the identification of Botrytis genes that were regulated in a host specific manner.

**Figure 1.**
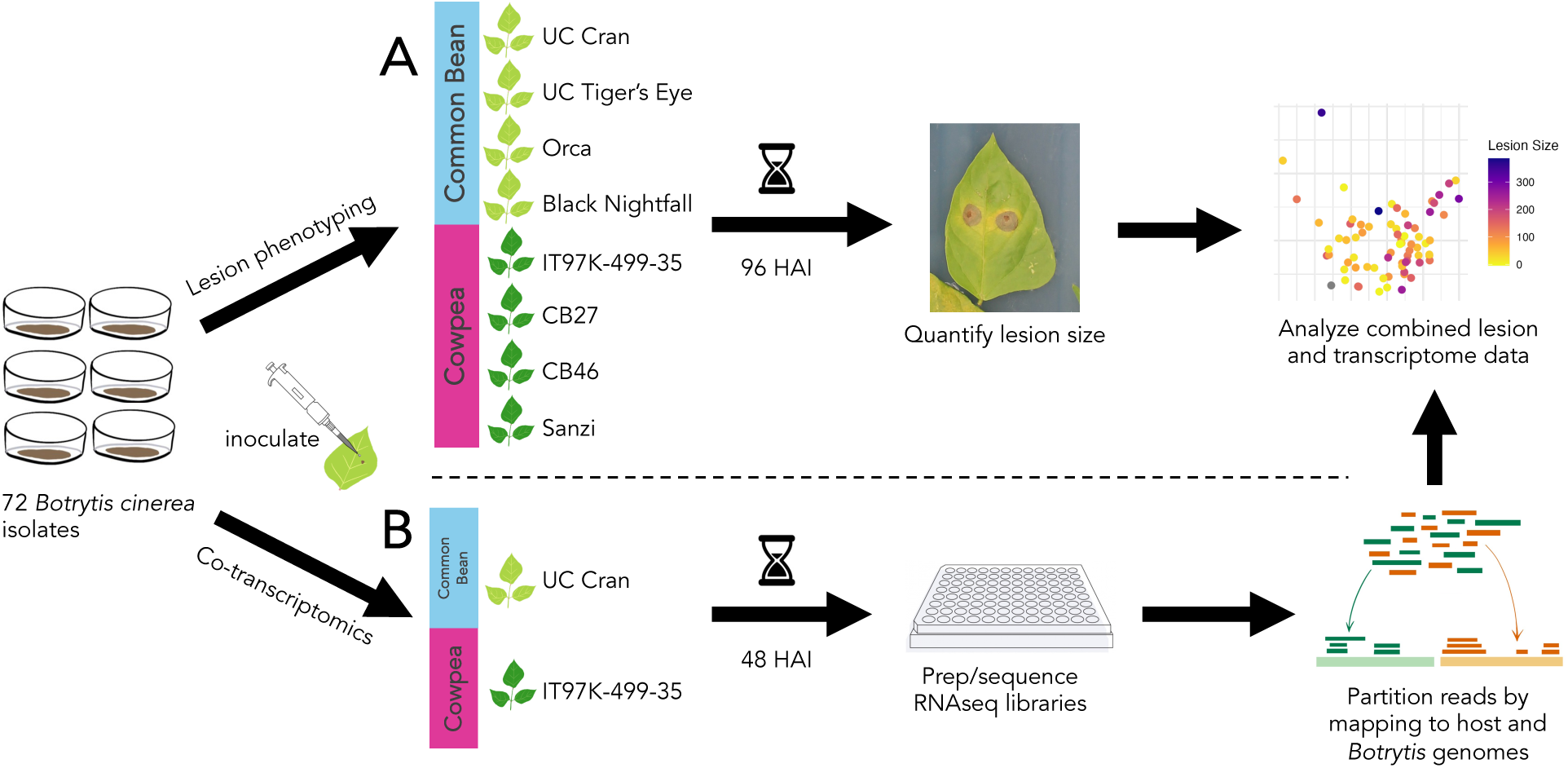
Schematic of experimental design. Lesion phenotyping was conducted across two independent experiments each with 3 biological replicates, and transcriptomics was conducted as one experiment with 3 biological replicates. HAI = Hours after inoculation.

## Results

### Lesion size and phenotypic variation

We measured Botrytis lesion formation on cowpea and common bean to test how the difference between closely related hosts may influence host-Botrytis interactions. 72 diverse Botrytis isolates were individually inoculated on leaves from 4 representative genotypes each of common bean and cowpea. Since different Botrytis isolates are a major driver of disease outcomes (Caseys et al. 2021; Singh et al. 2025), these isolates were used to assess what responses are stable in the host interactions with the pathogen versus what responses are conditional on pathogen variation. Disease phenotype data was collected in the form of digital images of lesion sizes for each combination of host-pathogen genotypes (Figure 1). Using a long-established assay for detached leaf assays (Denby et al. 2004; Caseys et al. 2021), the developing lesions were imaged at 24, 48, 72, and 96 hours after inoculation (HAI). Lesions became visible after 48 HAI, so lesion areas were digitally quantified only for 72 and 96 HAI timepoints. Lesions on common bean and cowpea grew quickly from 72 HAI to 96 HAI, but the rank order of the Botrytis isolate virulence between these timepoints was largely retained (Figure S1). To capture variation in lesion progression across isolates, lesion areas at 96 HAI were used for further analysis.

While cowpea showed slightly higher lesion size overall at 96 HAI compared to common bean, the genotypes within each host species showed a similar distribution of lesion sizes across the diverse Botrytis isolates (Figure 2A). This was reflected when conducting a linear model combining the lesion data across both host species. In this model, the terms host genotype, host species, and Botrytis isolate all significantly contribute to lesion size, where isolate was the strongest contributor (Table S1). In agreement with the strong isolate effect in the model, the lesion formation of the isolates was linearly correlated across the two species (Figure 2B). Interaction terms for host genotype x Botrytis isolate and host species x Botrytis isolate were not significant, indicating that we detected minimal variation of individual isolate’s lesion size across different hosts (Table S1, Figure 2C).

**Figure 2.**
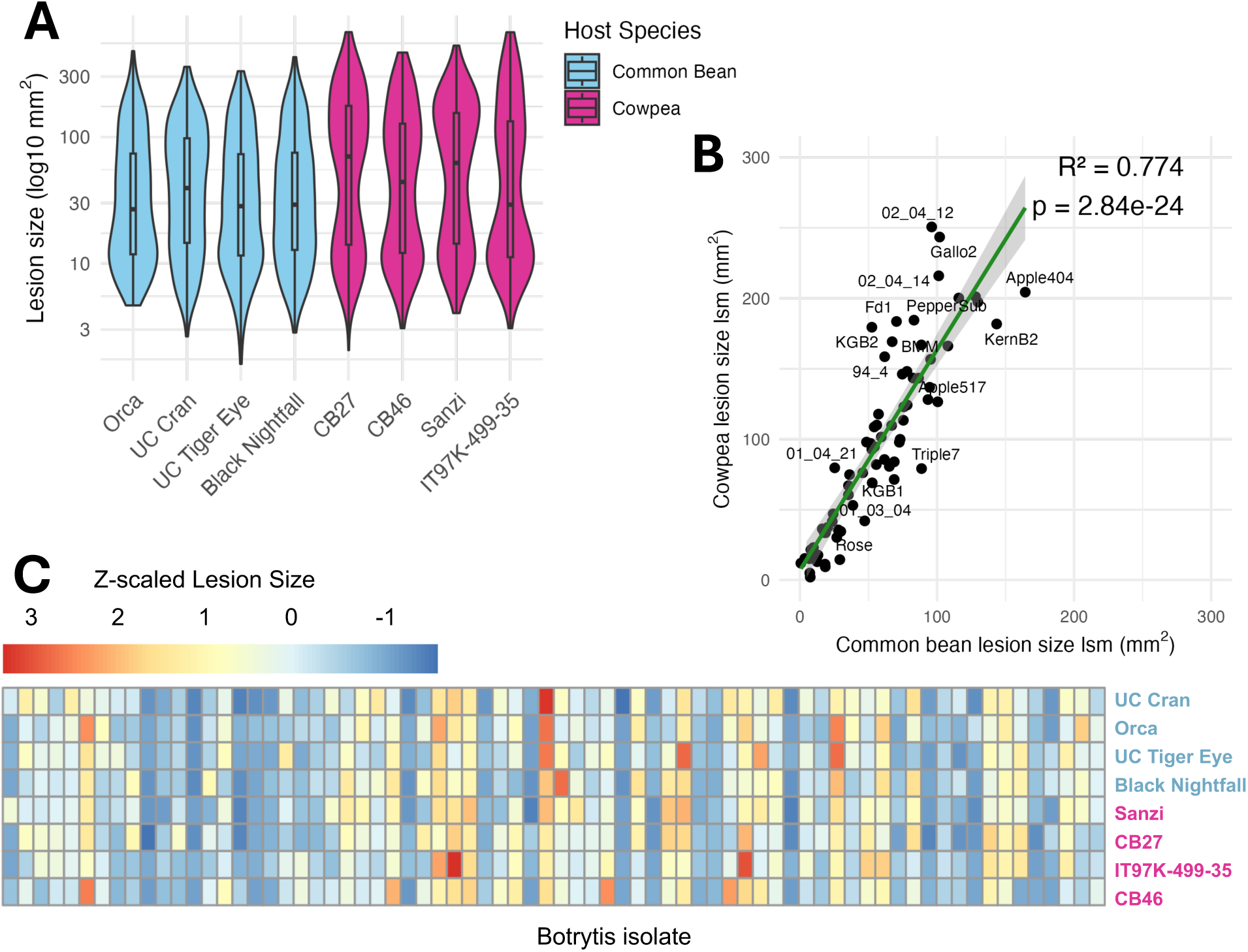
Lesion sizes of 72 *Botrytis* isolates on two legume host species at 96 HAI. A) Violin plot shows the overall distribution of lesion sizes collected for each host genotype. B) Correlation of host lesion size for each isolate. Species lesion size was calculated as the least-square mean (lsm) across the host genotypes. Outlying isolates are labeled with the isolate name. C) Lesion size heatmap showing z-scaled lesion sizes for each individual host genotype and Botrytis isolate. Common bean genotypes are colored in blue and cowpea genotypes are colored in magenta.

To quantify how the Botryis isolates’ lesion formation differs across host genotype variation for both host species, we calculated average species lesion residuals for each isolate (Figure S2). This was calculated for each isolate by subtracting the average lesion size for the isolate on a host species from its mean on each specific host genotype. These host genotype-level residual absolute values were then summed to generate one host species-level residual value for each host species and isolate combination. Therefore, an isolate with a residual value near 0 has low within-species specificity, while an isolate with a residual value far from 0 has high within-species specificity. This showed that several isolates are more sensitive to within-host species genetic variation, but the specific Botrytis isolates that display this host genotype dependency are independent across the two host species. Together, this suggests that host-Botrytis interactions in these two closely related legumes are largely driven first by variation in the pathogen and secondly by variation in the two host species.

### Botrytis transcriptome differs across two legume host species

Since the observed disease outcomes between closely related hosts were similar to each other, we wanted to test if the underlying transcriptional machinery governing this outcome is also similar between hosts. To test this, we next proceeded to measure the transcriptomic responses of the 72 Botrytis isolates against cowpea and common bean. The co-transcriptome was obtained by extracting total RNA from the developing lesion at 48 HAI and sequencing both the host and pathogen transcriptomes from the same samples and libraries (Figure 1). The following section focuses solely on analyzing the pathogen’s transcriptome.

Principal component analysis (PCA) showed that the overall Botrytis transcriptome had a difference in response to the two host legumes (Figure 3A). Comparing these transcriptome vectors to lesion formation in the two hosts showed that the main axis of Botrytis isolate transcriptome variation (PC1) does not significantly associate with lesion size (p = 0.22), but the second axis (PC2) does significantly associate to lesion size (p = 0.003). However, PC2 is only 3.67% of the overall variance in the Botrytis transcriptomic dataset, suggesting that Botrytis gene expression is not the only contributing factor to disease outcomes. PC2 also significantly associates with the total Botrytis transcript abundance (total number of mapped Botrytis reads) for the sample (p < 0.001) (Figure S3). This analysis shows that while Botrytis transcriptomic variation between hosts was evident, only a small proportion of this variation was significantly associated with lesion size and overall transcript abundance, and likely other biological or environmental factors also influence transcriptomic variation.

**Figure 3.**
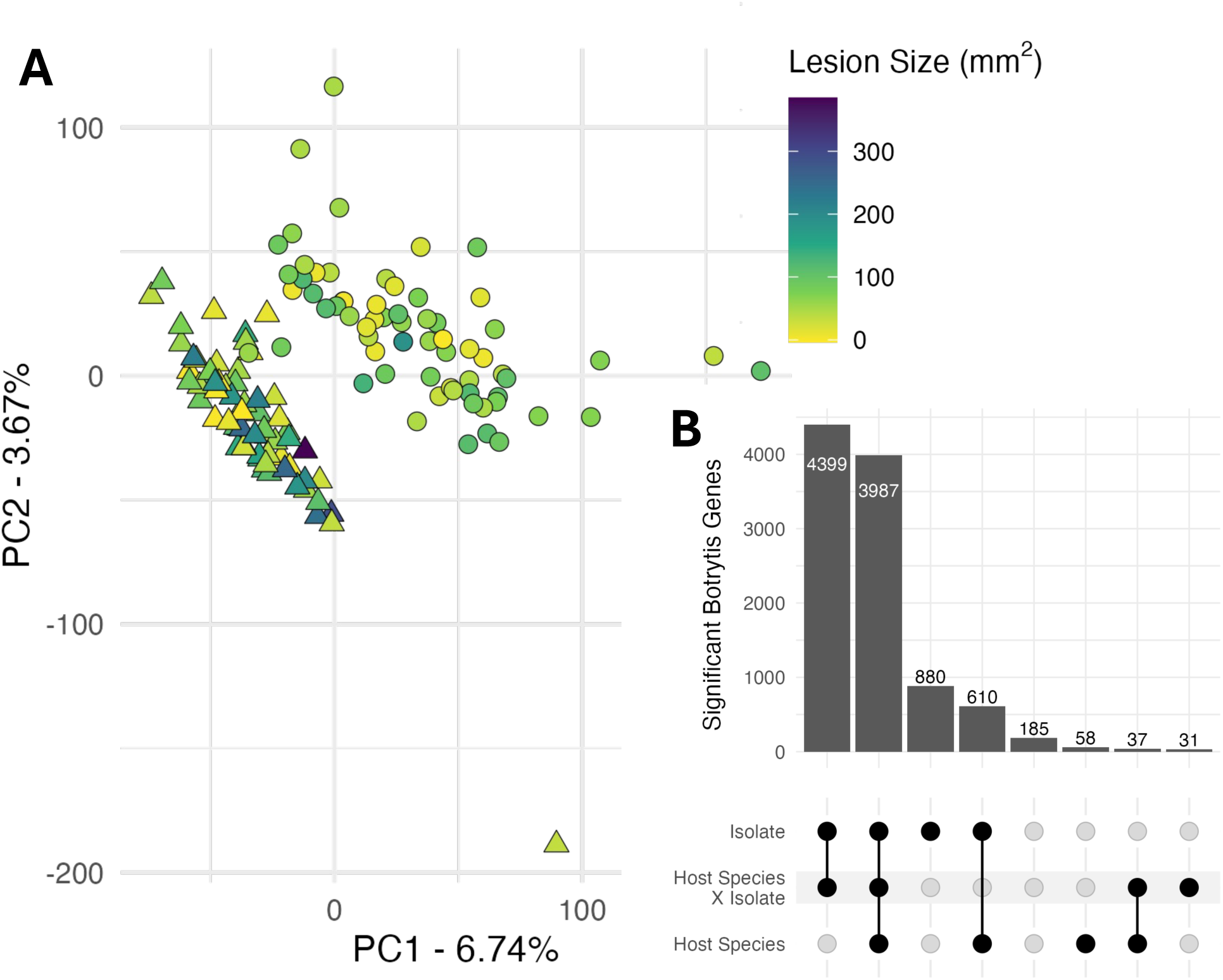
Overall Botrytis transcriptome variability during infection of two legume host species. A) PCA of overall Botrytis transcriptome at 48 HAI. Shape of points show the species the infecting Botrytis isolate was collected from, where circles are common bean and triangles are cowpea. Points are colored by lesion size of that Botrytis isolate on that host species at 96 HAI. B) Upset plot summary of ANOVA for each Botrytis gene expression on legume hosts at 48 HAI. Botrytis genes were separately modeled with the formula: gene expression ∼ host species + isolate + (host species * isolate). Genes are included in each category if the FDR < 0.05 for that model term. Genes with little to no expression are not diagrammed and include 1,887 Botrytis genes.

Moving beyond the whole transcriptome level to the individual transcript level, 10,187 out of 12,111 total genes in the Botrytis genome could be quantified after quality control and low readcount filtering. The vast majority of Botrytis genes (83%) were differentially expressed during infection, either between the host species, amongst the Botrytis isolates, or across the interaction. Assessing by the number of genes associated with each term, Botrytis gene expression variation across the dataset was first influenced by the Botrytis isolate, secondarily by host species x Botrytis isolate interaction, then thirdly by host species (Figure 3B). This result contrasts with the lesion modeling, which showed little to no host x isolate interactions in lesion formation. With a large portion of genes having a host x isolate effect on their expression, this suggests a large amount of variation of underlying host x isolate effects on gene expression was masked beneath the lesion phenotype.

### Differential expression of Botrytis genes across hosts highlights host-specific virulence mechanisms

We hypothesized that the Botrytis genes showing a large difference in expression between common bean and cowpea could provide insight into how Botrytis’s virulence strategy might change across these closely related hosts. Of the 4692 Botrytis genes with a significant host effect, only 58 Botrytis genes had solely a host effect with no isolate variation. GO enrichment of these 58 genes against the whole Botrytis genome showed enrichment in protein binding and endoribonuclease activity (Table S2). To look more broadly at host specific Botrytis genes, we focused on all 4692 genes with a host effect regardless of whether they also had an isolate effect, as this offered a more inclusive assessment of the differences in response to the hosts. Filtering the 4692 total Botrytis genes with a host species effect for those with at least a two-fold difference between the two hosts yielded 1078 differentially expressed Botrytis genes (Table S3). This included 656 Botrytis genes significantly upregulated during infection of cowpea in comparison to common bean, and 422 Botrytis genes significantly upregulated during infection of common bean in comparison to cowpea (Figure 4A). Among these, many of the most differentially expressed genes are in gene families known to aid in cell wall penetration and host surface modification. Several identified genes encode such functions like glycosyl hydrolases, glycosyl transferases, cellulose degrading, cutinases, etc. This suggests that Botrytis may be responding to common bean and cowpea differently by altering cell wall-degrading gene expression. In addition to cell wall metabolism, a number of the genes are involved in unknown specialized metabolite pathways, including polyketide synthases and cytochrome P450s.

**Figure 4.**
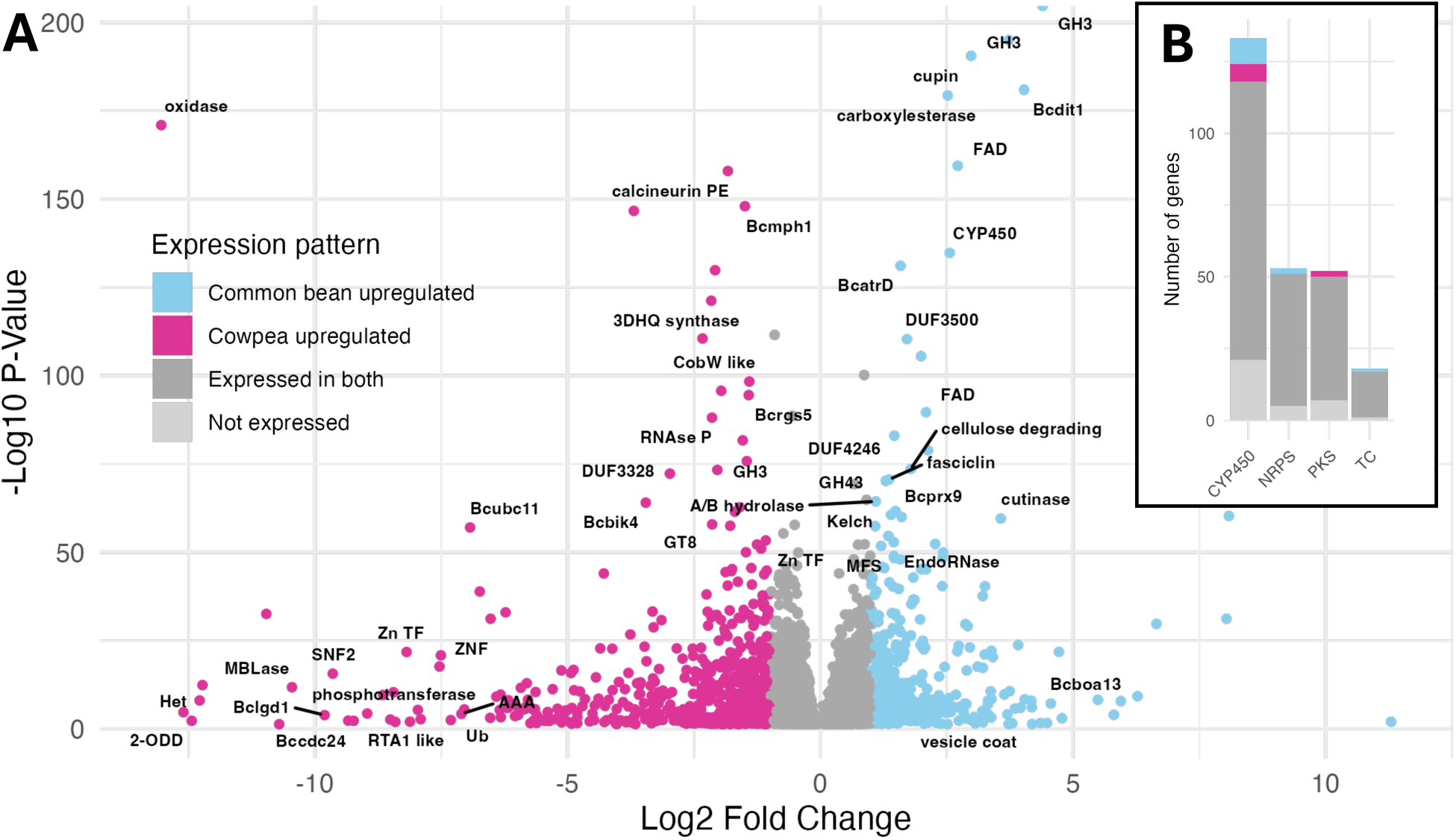
Host-specific expression of Botrytis genes. A) Volcano plot of Botrytis genes plotted by the gene’s log2 fold change value when infecting common bean vs. cowpea. Annotated genes with -Log10 P-value > 50, log2FC > 4, or log2FC < -7 are labeled on the plot. B) Count of genes in key secondary metabolic gene classes for Botrytis, colored by their host specific expression patterns. CYP450 = cytochrome P450, NRPS = nonribosomal peptide synthase, PKS = polyketide synthase, TC = terpene cyclase.

To measure the relative host specificity of Botrytis specialized metabolic gene expression, we used a previously described annotation of known specialized metabolic enzymes in Botrytis (Suárez et al. 2024). This allowed for classification of enzymes into 4 major categories: cytochrome P450s (CYP450), nonribosomal peptide synthases (NRPS), polyketide synthase (PKS), and terpene cyclases (TC), including both diterpene and sesquiterpene cyclases. All genes in each specialized metabolic class were totaled within their observed regulatory pattern groups on the hosts (up on common bean, up on cowpea, up on both hosts, or up on neither host). While most of these specialized metabolic genes were expressed when infecting both hosts, a small number of genes in each class showed host specificity (Figure 4B). This indicates Botrytis displays small adjustments in specialized metabolic pathways even when infecting closely related hosts. Each regulatory pattern gene group (upregulated in common bean only, upregulated in cowpea only, upregulated in both, or upregulated in neither) was then tested for enrichment of these 4 specialized metabolic classes against the ratios of specialized metabolic genes present in the whole genome using the hypergeometric distribution. No single specialized metabolite class was significantly enriched during infection of either host, indicating that Botrytis uses a blend of specialized metabolites depending on the host overall.

### Key Botrytis phytotoxin pathways express similarly on legume hosts across diverse isolates

In addition to differences between the hosts, genetic variation between the 72 Botrytis isolates was a major driver of lesion and Botrytis transcriptome variation (Figure 3B). To better understand which Botrytis virulence mechanisms may be affected by isolate variation, yet likely function equally across the hosts, we generated a list of the 880 Botrytis genes with a significant isolate effect and no significant host associated variation. This list represents genes that vary in expression across isolates, but are expressed similarly when infecting either host species. To obtain a clearer mechanistic signal from this large list of genes, Botrytis gene co-expression networks (GCNs) were generated separately using each of the 2 host datasets using all expressed genes. These GCNs were filtered for those where at least 25% of genes in the GCN are from the list of 880 Botrytis isolate-specific genes to identify common Botrytis GCNs expressed similarly across hosts, but altered by the pathogen’s genetic variation. These identified 18 isolate-specific Botrytis GCNs, including 10 found when infecting common bean and 8 when infecting cowpea (Table S4). Among these networks were one including BcVEL1, a gene from the VELVET regulatory complex known to be genetically variable and influence virulence across hosts (Schumacher et al. 2012).

In this list of similarly expressed GCNs across hosts, there were also Botrytis GCNs that included the well-described biosynthetic pathways for key phytotoxins, botrydial and botcinic acid. These biosynthetic pathways exist as two gene clusters each co-localized on separate chromosomes (Table S5). The expression of these phytotoxin clusters exhibits a quantitative and qualitative variation across the isolates, where most isolates express the phytotoxins on both hosts, and several isolates have very low or no expression of one or the other pathway on either host (Figure 5). This suggests Botrytis expresses botrydial and botcinic acid similarly across both closely related hosts, and these virulence mechanisms may be an example of a generalized defense. Still, individual isolates varied widely in their expression of these pathways, and the variation in the two phytotoxins was independent such that there are a few isolates that have lost or dramatically decreased the expression of one cluster or the other on a specific host. However, the isolates that have low expression of these phytotoxin pathways overall do not have apparently reduced virulence (Table S6) and are likely relying on other mechanisms to induce host toxicity, consistent with previously reported functional redundancy of these phytotoxins (Dalmais et al. 2011; Leisen et al. 2022).

**Figure 5.**
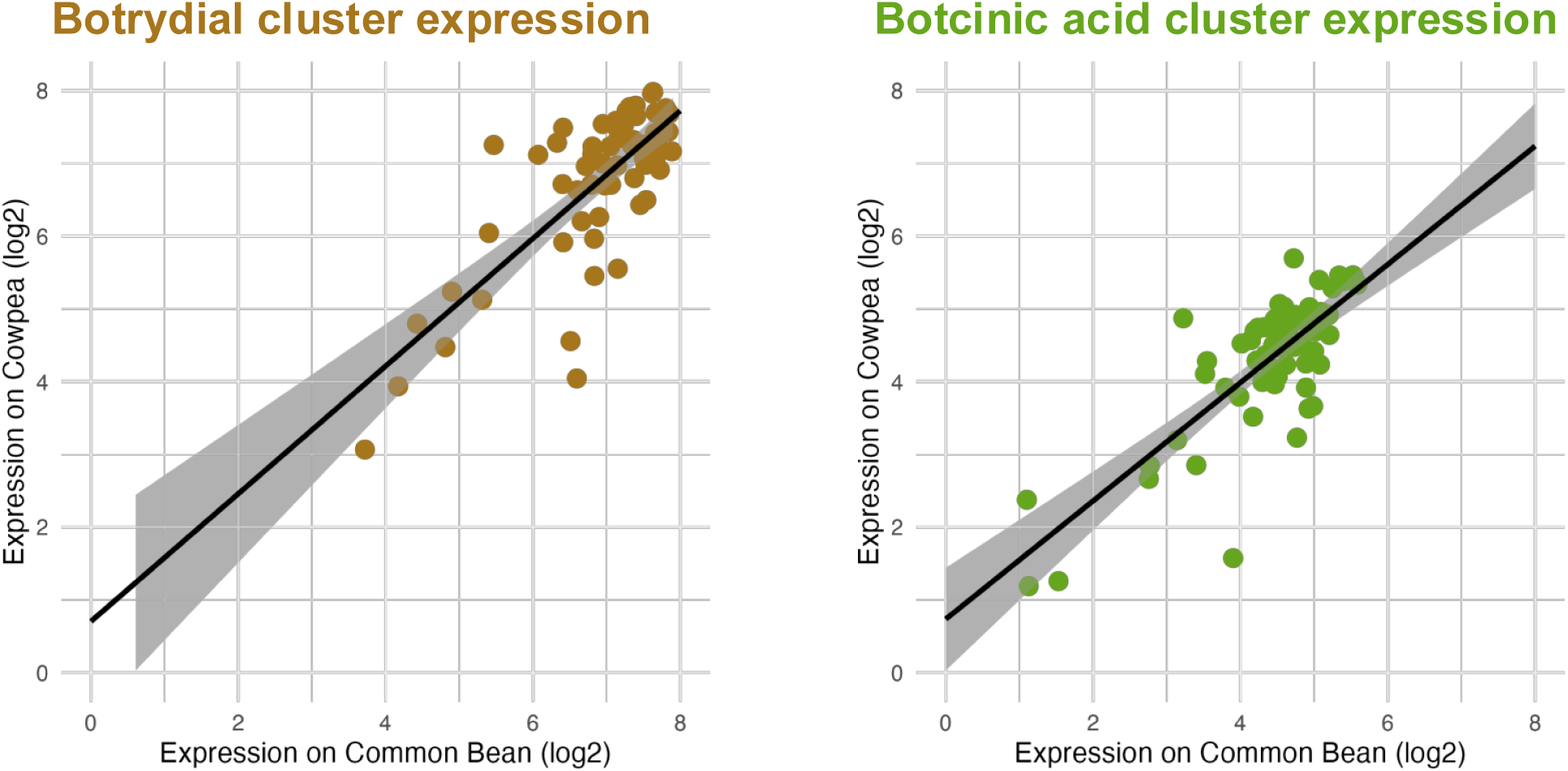
Expression of Botrytis phytotoxin biosynthetic pathways across host species. Average z-scaled expression (log2) was calculated for known genes in each pathway for each Botrytis isolate. Host-specific expression of each isolate was plotted. Botrydial pathway consisted of a 5 gene cluster (Bcin12g06370-Bcin12g06410), while botcinic acid pathway consisted of an 16 gene cluster (Bcin01g00010-Bcin01g00160).

### Botrytis gene co-expression networks vary across host and isolate genetic diversity

Most Botrytis genes’ expression have a significant host x isolate interaction, meaning their expression varies across the hosts depending on which isolate is infecting the hosts. The observation that host x isolate interactions had minimal influence on lesions but large effects on the transcriptome suggested two possible models. First, each host induces different Botrytis networks that are variable across isolates and sum to create similar lesion variation. In the second model, the host effects reshape similar networks across the isolates leading to similar lesions. To test these models, we assessed if the host x isolate Botrytis GCNs found on cowpea and common bean were different (Model 1) or contain subsets of same genes (Model 2). This network approach can identify mechanisms influenced by these host x isolate interactions and provides insights on how Botrytis infection mechanisms differ across genetic diversity in both the host and the pathogen.

To identify the different Botrytis mechanisms influenced by the host x isolate interaction, the full set of Botrytis GCNs were filtered to identify networks that contained at least 5 of the top 100 host x isolate interaction-significant Botrytis genes based on p-value of the interaction term. This approach found 4 Botrytis GCNs when infecting common bean and 5 when infecting cowpea. Because the Botrytis GCNs represent infection of two different host datasets, GCNs that contain similar sets of genes can be identified on either host. Among this set of 9 Botrytis GCNs were 2 pairs of similar GCNs across the hosts. The first similar Botrytis GCN found on both host species identifies a co-expressed and previously undescribed gene cluster on chromosome 13 that may be responsible for breakdown and potentially detoxification of host compounds (Figure S4). The second similar Botrytis GCN centers around a nonribosomal peptide synthase (NRPS) gene cluster on chromosome 12 (Figure 6A & B). Comparing z-scaled expression values of these networks for each host-isolate combination shows large differences in network expression across hosts for several isolates (Figure 6C). In addition to differences in expression on the hosts, gene memberships in the networks can shift. While these 2 GCNs have some identical gene members shared across both hosts, there are key differences in overall network structure when infecting common bean vs. cowpea. Interestingly, when infecting common bean, the NRPS cluster co-expresses with several shikimate pathway genes (EPSP synthase, BcPHA2, BcARO2 and BcARO7) as well as an additional polyketide synthase (PKS) gene cluster from chromosome 14 (Figure 6A). Many fungi have been shown to have hybrid PKS-NRPS enzymes that catalyze steps in flavonoid biosynthesis, and can do so directly from p-coumaric acid (Zhang et al. 2022). However, when infecting cowpea, the Botrytis PKS gene cluster GCN lacks co-expression with these shikimate related enzymes. Instead, the network is modified such that the NRPS cluster co-expresses most closely with several tailoring enzymes that could modify a NRPS-based metabolite, including an O-methyltransferase, amidohydrolase, laccases (CLCC2 & CLCC7), and dehydratases (Figure 6B). This creates a system whereby the core PKS-NRPS module seems to have different potential tailoring/modification genes depending on the host and the isolate. These modified NRPS networks provide an example of a Botrytis response system potentially encoding unknown branching metabolic pathways that can be observed across genetically diverse interactions, but differentially fine-tuned depending on both the host genotype and the specific Botrytis isolate. This supports the second model proposed above: closely related hosts reshape similar Botrytis networks across the isolates, but these changes lead to similar lesion variation.

**Figure 6.**
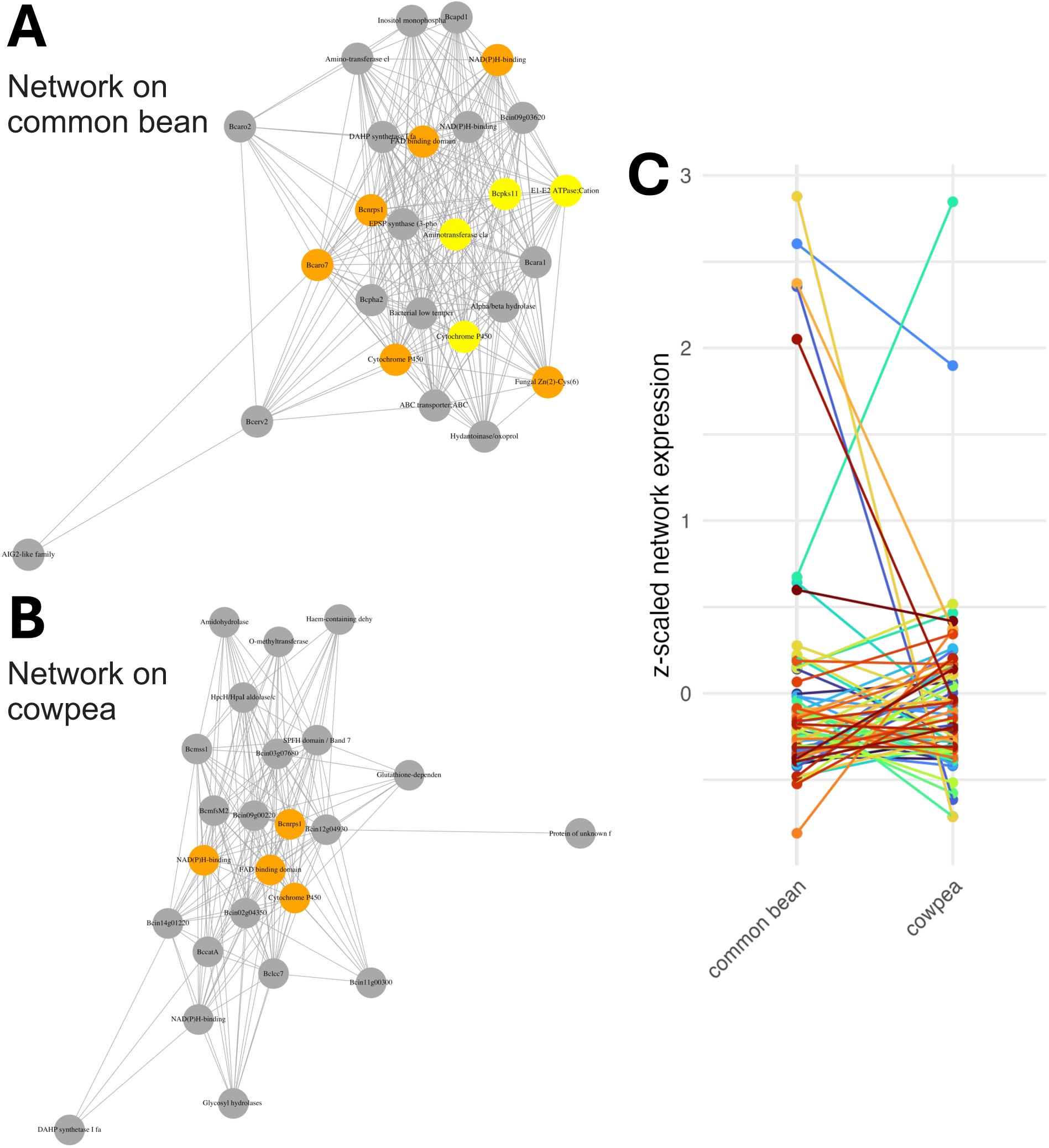
Selected Botrytis co-expression networks containing several genes with significant host x isolate interaction. Similar Botrytis gene co-expression networks were detected when infecting both A) common bean and B) cowpea with slight differences. Biosynthetic gene clusters detected within each network are highlighted in color, where the NRPS cluster on chromosome 12 is in orange and the PKS cluster on chromosome 14 is in yellow. C) Z-scaled expression of these interaction networks on different hosts. Colored points and connecting lines represent different Botrytis isolates.

### Host species transcriptomes response to Botrytis infection is shaped by Botrytis genetic diversity

The co-transcriptome approach allows us to simultaneously measure the hosts’ gene expression in these exact same samples and compare and contrast the host’s response with the pathogen’s transcriptome. For each host species, there was a single genotype used and time-matched uninfected samples were included as controls. Each host gene’s expression was modeled with the formula: *gene expression ∼ infected + infected/isolate.* In contrast to the Botrytis model, this had to be run independently on the two hosts as the genes are different in the two species. The proportion of genes that responded significantly to infection was considerably higher in common bean (26%) than in cowpea (12%). However, in both hosts, a large proportion of host genes were differentially expressed in response to the diverse isolates, 42% for common bean and 37% for cowpea (Figure 7). This is in agreement with variation in Botrytis isolate being a key determinant of lesion formation in both hosts. A key difference between the two host models is that a higher proportion of genes show an infection effect in common bean than in cowpea. This suggests that although overall variance in host gene expression across isolates is similar between the two species, cowpea exhibits a weaker average transcriptional response to Botrytis than common bean. This may partially explain the slightly larger average lesion size for cowpea (Figure 2A).

**Figure 7.**
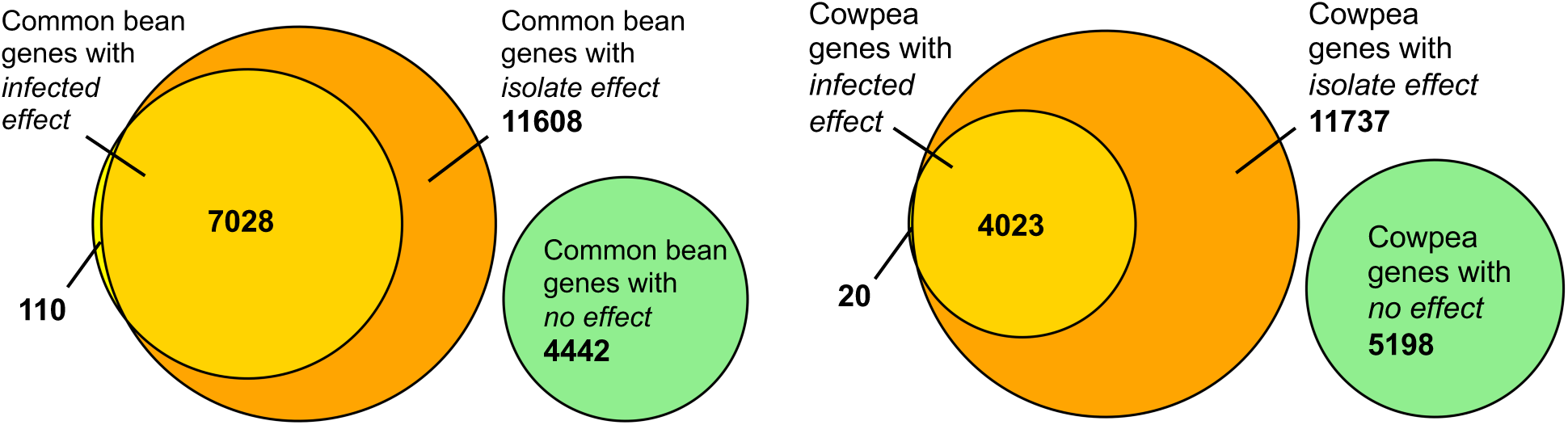
Summary of ANOVA for each legume host gene expression at 48 HAI with Botrytis. Host genes were separately modeled with the formula gene expression ∼ infected + infected/isolate. Genes are included in each category if FDR < 0.05 for that model term. Genes with little to no expression are not diagrammed and include 11,273 genes for common bean and 14,993 genes for cowpea.

To assess the overall behavior of the two host transcriptomes, we initially conducted PCAs that showed the PC1 in common bean explained 39% of its transcriptomic variation, while the PC1 in cowpea explained 42% of its transcriptomic variation (Figure S5). In both hosts, PC1 associates with the estimated Botrytis biomass (as measured by total reads mapped to the pathogen; Botrytis transcript abundance) (Figure S5A). Interestingly, PC1 in either host does not associate with lesion size at 96 HAI (Figure S5B). Because host gene expression is not strongly associated with disease incidence but is instead more responsive to the abundance of Botrytis transcripts this suggests that host response during early infection is influenced primarily by variation in the Botrytis isolate. This agrees with previous observations that lesion formation and Botrytis biomass are only mildly correlated (Corwin, Subedy, et al. 2016).

### Host orthology analysis reveals several conserved patterns in response to Botrytis

To enable a direct comparison of host responses between the two host species, we conducted a genome wide orthology analysis to procure a list of single copy orthologs between common bean and cowpea. This identified 18,913 genes that are single copy orthologs between common bean and cowpea. These are genes that have a single gene match across the legume species and allow us to directly compare the transcriptome response to Botrytis between both host species. A majority (∼70%) of the expressed host genes in the study comprise the list of single copy orthologs, a representative subset of both the cowpea and common bean genome (Figure S6).

Of the 18,913 single copy orthologs, 2,923 were expressed in common bean but not cowpea, 2,509 were expressed in cowpea but not common bean, and 5,294 were not expressed in either host. The remaining 8,187 single copy orthologs expressed in both hosts were then filtered to the 2,541 that had a significant response to Botrytis infection in both hosts. Using this list of orthologs, we directly compared the two host species’ transcriptomic responses to Botrytis infection by calculating the mean expression of each gene in the mock sample and across all the Botrytis isolates. This was used to calculate the log2 fold change (log2FC) of each transcript in response to infection within each host species. Plotting the log2FC values of each single copy ortholog across the two host species allowed for splitting the genes into 4 quadrants: upregulated in both hosts, downregulated in both hosts, upregulated in common bean but downregulated in cowpea, and downregulated in common bean but upregulated in cowpea (Figure 8, Table S7). This showed that while most host orthologs show similar responses to infection, there is a set of genes with opposite responses to infection in the two host species.

**Figure 8.**
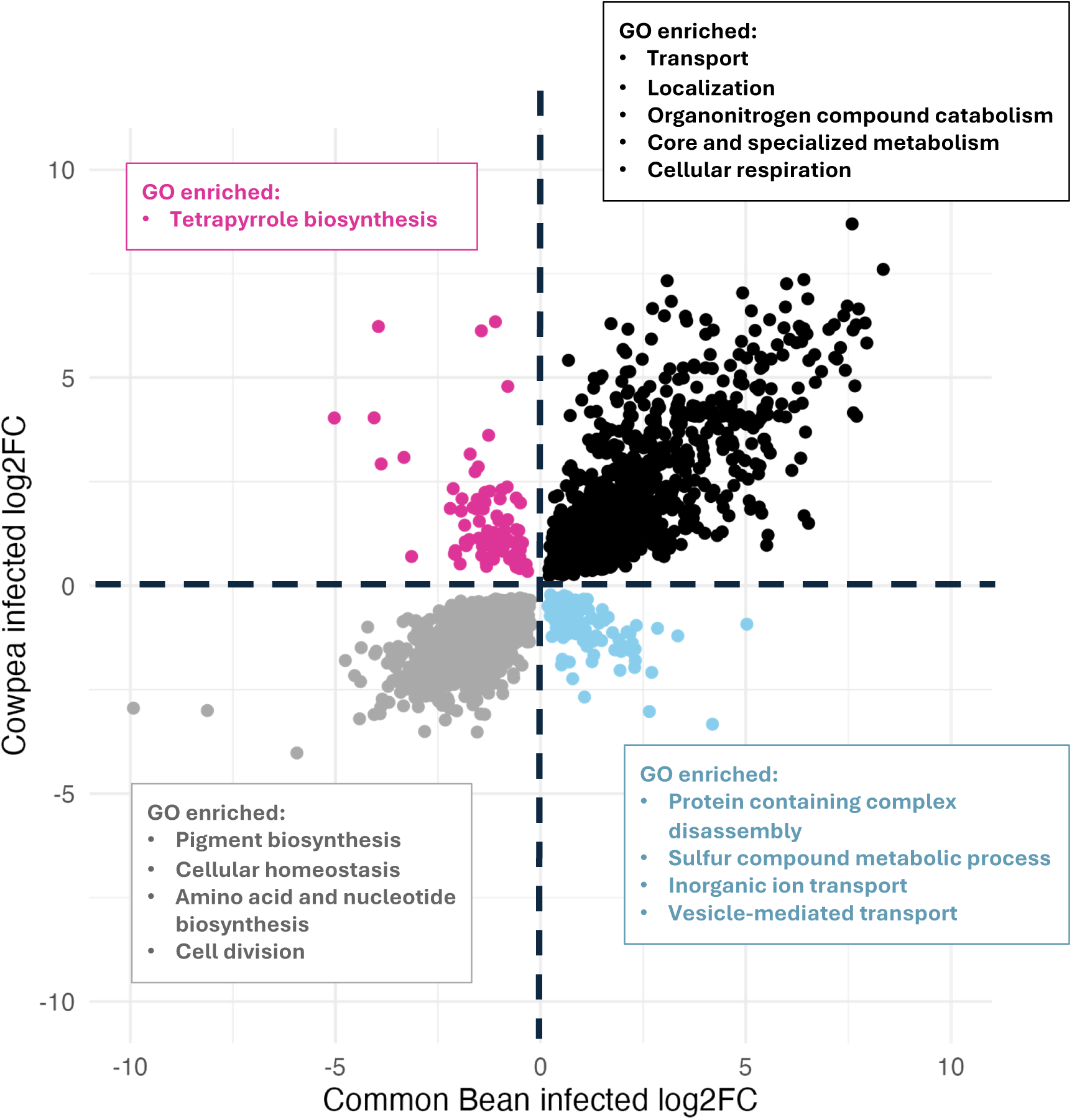
Orthology reveals shared and host-specific responses to Botrytis. Comparative log2FC of single copy orthologs between cowpea and common bean that have a significant response to infection in both host species. Log2FC is calculated as the difference of expression from control to infected. Genes are colored by their expression pattern, where magenta is upregulated in cowpea and downregulated in common bean, blue is upregulated in common bean and downregulated in cowpea, black is upregulated in both, and grey is downregulated in both. Summary of significantly enriched GO terms (p < 0.05) are shown for each quadrant. Only genes significantly altered by infection are shown (FDR < 0.05).

GO analysis on the 1235 genes upregulated by Botrytis infection in both host species identified 86 significantly enriched GO terms. The most significant terms were dominated by those related to localization and transport, but many significant terms also included cellular respiration and core cellular metabolic processes, including terms for mono- and dicarboxylic acids, pyruvate, nucleotides, etc. (Table S7). Additionally, several pathways producing specialized metabolite precursors were upregulated in both hosts including malonate and isoprenoid biosynthesis. These pathways potentially feed into the downstream production of flavonoids and terpenes. Defense signaling pathways were also upregulated in both hosts, including jasmonic acid pathway genes (JAZs, LOX3, OPR3) and ethylene pathway genes (ACC synthase, ERF1).

Testing the 1088 single copy orthologs downregulated in both hosts during infection identified 39 GO terms. Overall, these terms were more representative of processes associated with photosynthesis and growth, including pigment, amino acid, and nucleotide biosynthesis, cell division, and cellular homeostasis. These processes are also downregulated in other diverse host plants when infected by Botrytis (Windram et al. 2012; Zhang et al. 2017; Zhang et al. 2019; De Cremer et al. 2013).

There were 218 single copy orthologs differentially responsive in the two species (Figure 8). Relatively few genes were downregulated in common bean but upregulated in cowpea, with only 2 enriched GO terms both relating to tetrapyrrole biosynthesis. For those genes upregulated in common bean but downregulated in cowpea, these were enriched for 5 GO terms, including complex disassembly, sulfur compound metabolism, and transport.

### Co-expression analysis reveals differences in upregulated host responses to Botrytis

To provide more information on the processes represented in the single copy orthologs in the genomic contexts of both hosts, we used the host GCNs. The host GCNs were determined separately for each host species so that networks could be built with both orthologous and non-orthologous genes within each host. The networks were filtered to those that had at least 10 members, and had at least 50% of its members as a host single copy ortholog significantly altered by infection in both hosts (same genes as Figure 8). This resulted in 37 infection-responsive host networks, 20 for common bean and 17 for cowpea. Approximately half of these networks contained orthologs that were downregulated in both hosts, mainly annotated as photosynthetic genes (Table S8).

To look more closely at processes upregulated in response to Botrytis in both hosts, we focused on the upregulated networks, which included 10 for common bean and 8 for cowpea. These networks contained a blend of single copy orthologs and other genes specific to each host (Figure 9A). Since the majority of the network members were single copy orthologs upregulated in response to Botrytis in both hosts, we hypothesized the genes are similarly regulated in either host. Therefore, we expected that corresponding networks across hosts would share a high fraction of genes. However, we found a relatively low degree of similarity in gene content between the upregulated host networks (Figure 9B). The 5 most similar networks (ranging from 14-35% match) all resembled various transport/signaling pathways, with the exception of two biosynthetic networks (PV474 and VU192). These two networks were chosen as an example to compare similar responsive pathways between the two hosts (Figure 9C). The shared single copy orthologs between these networks are 3 genes: phenylpropanoid pathway gene C4H (cinnamate-4-hydroxylase), aquaporin protein SIP1, and acyl-activating enzyme 18 (AAE18). The other network genes have connections to shikimate and phenylpropanoid metabolism, but are different specific genes in each species. This shows shikimate and phenylpropanoid metabolism, while upregulated in both common bean and cowpea, have specific differences in response to Botrytis infection. Thus, even for shared upregulated networks in closely related hosts, the hosts differ in specific network composition.

**Figure 9.**
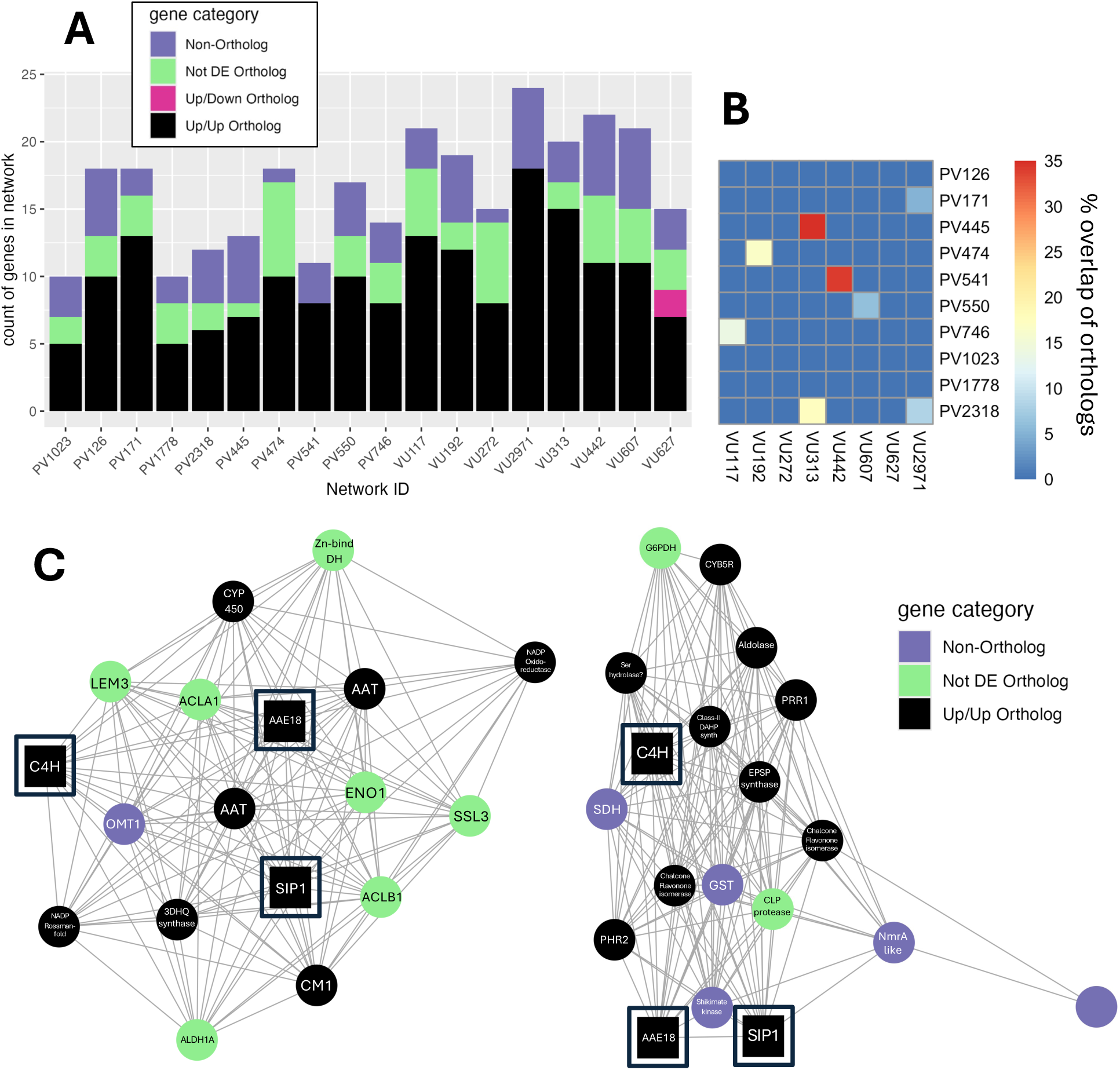
Upregulated host networks share similarities in overall function, but high variability in gene membership across host species. A) Total genes present in each upregulated host network, colored by orthology and regulatory category. Up/Up ortholog = upregulated in both hosts during infection, Up/Down ortholog = upregulated in one host and downregulated in the other during infection, Not DE ortholog = not differentially expressed during infection, and Non-Ortholog = a gene without single copy orthology between the hosts. B) Heatmap showing network homology via the percent of overlap among single copy orthologs across networks. C) Upregulated phenylpropanoid biosynthetic GCNs from common bean (PV474) and cowpea (VU192) with homology across hosts. Squares represent single copy orthologs with overlap across the networks.

Some of the upregulated single copy orthologs in Figure 8 were from the well known hormone defense-signaling pathways jasmonic acid (JA) and ethylene. These pathways have previously been shown to be involved in plant defenses against Botrytis. Therefore, we specifically investigated upregulated host GCNs for the presence of JA or ethylene networks. There were 4 single copy jasmonate-ZIM domain orthologs that were upregulated in both hosts. Of these, none appeared in co-expression networks for common bean. 2 of them, JAZ1 and JAZ6, appeared across 3 GCNs in cowpea. Interestingly, these networks did not include other known and upregulated JA pathway genes (i.e. LOX3, OPR3). For ethylene signaling, there were 3 ERF1 orthologs upregulated in both hosts. Of these, none appeared in co-expression networks for cowpea. One appeared in a network for common bean (PV171), otherwise this network’s known members consist of those involved in signaling and transport (Table S8). This suggests that these defense response networks are differentiated between common bean and cowpea.

### Most host orthologs respond differently to Botrytis isolates in either host

The composition of the host GCNs upregulated in both hosts suggests that while many of the same orthologs are generally upregulated during infection in either host, they co-express differently across hosts, likely depending on both the host and the infecting Botrytis isolate. We tested for these effects by modeling all single copy orthologs with the following model: Single copy ortholog expression ∼ host + isolate + host X isolate. This revealed that single copy genes shared between the hosts largely shift in their expression not only in either host, but also in response to genetic diversity in Botrytis. A large proportion of single copy orthologs had significant effects across host species, Botrytis isolate, and the interaction (Figure 10). The vast majority of the expressed orthologs (94%) were differentially expressed between host species, a result expected given that this analysis integrates gene expression from two distinct genomes. Despite this strong effect, isolate and host x isolate interaction also contributed substantially to host ortholog expression. Specifically, 81% of the plant orthologs were differentially expressed across isolates, indicating that pathogen genetic diversity broadly shapes host transcriptional responses and this differs between the two hosts. Orthologs that varied across isolates but not the interaction term likely represent responses relatively conserved between hosts. However, over half of host orthologs (53%) had significant host x isolate interactions, indicating isolate-specific effects that differ between hosts. Together, this shows that although these closely related host species share much of their genome, their shared gene expression programs are intricately fine-tuned in response to genetic diversity in a generalist pathogen.

**Figure 10.**
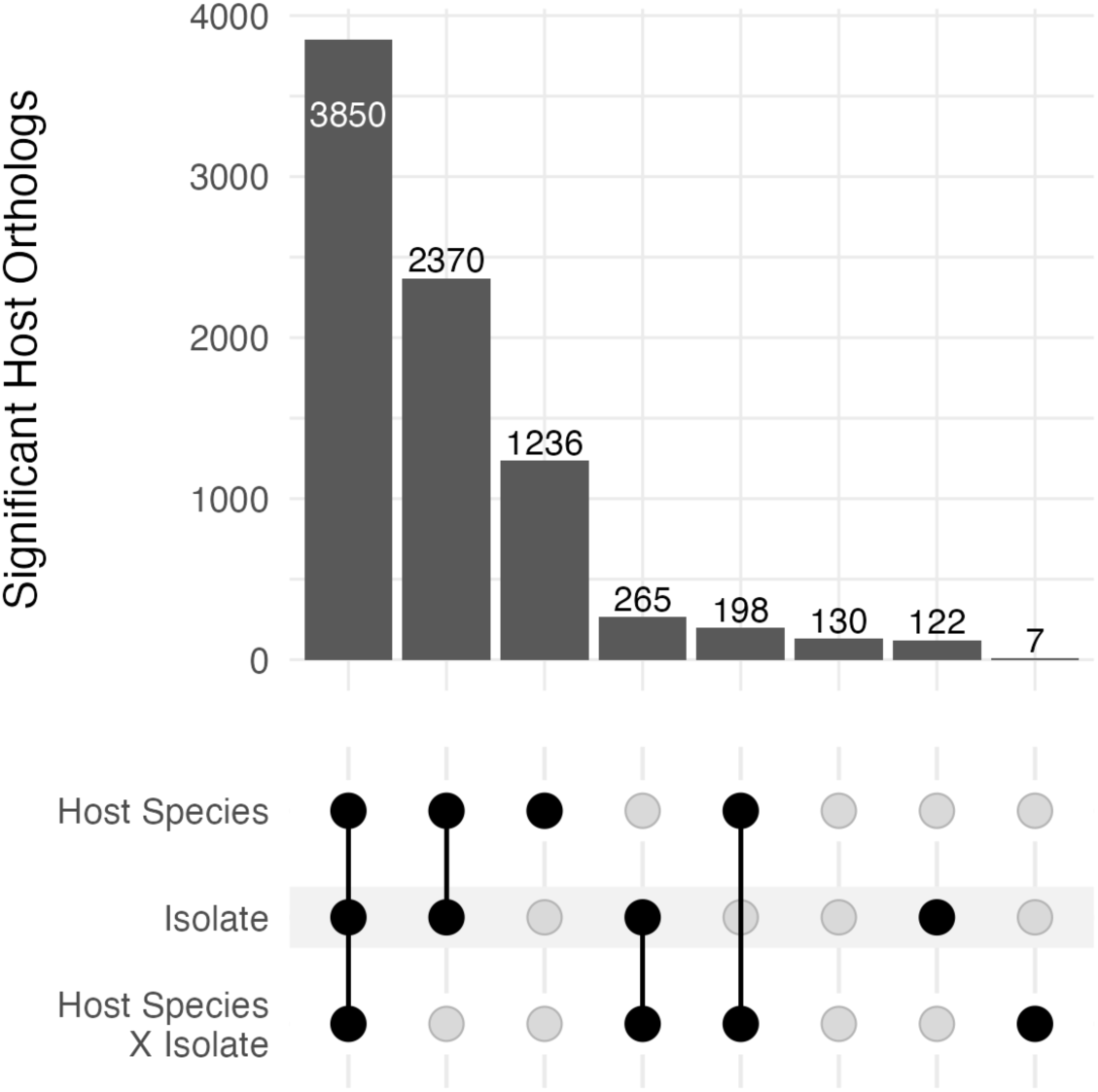
Most host single copy orthologs show differential expression across hosts and in response to Botrytis. Upset plot summary of ANOVA for each host single copy ortholog gene expression for legume hosts at 48 HAI. Host orthologs were modeled with the formula: Ortholog expression ∼ Host Species + Isolate + Host Species * Isolate. Genes are included in each category if the FDR adjusted p value was < 0.05 for that model term. Genes with little to no expression are not diagrammed and include 5,294 host orthologs.

## Discussion

To understand how differences between closely related host species shapes interactions with a generalist pathogen, we examined infection outcomes and co-transcriptomic responses of *Botrytis cinerea* and two closely related legume hosts, common bean and cowpea. Using a diverse panel of 72 Botrytis isolates, lesion formation and gene expression were quantified in both host and pathogen to separate the effects of host genetic diversity, pathogen diversity, and their interaction (Figure 1). This identified complex, fine-tuned molecular responses in both host and pathogen that were masked at the phenotypic level.

### Botrytis infection of closely related hosts reveals conserved and divergent molecular interactions

A central question in host generalism is how differences between host species shape molecular interactions during infection. For closely related hosts, this question narrows to 1) whether a generalist pathogen uses similar virulence strategies across hosts, and 2) whether related hosts respond similarly to the same pathogen. Recent work has shown that Botrytis displays host-specific gene expression across eudicots while maintaining a core of genes expressed across all hosts (Singh et al. 2025). Consistent with this, we observed a mixture of conserved and divergent responses in both host and pathogen transcriptomes, revealing substantial molecular divergence despite close host genetic similarity.

The Botrytis transcriptomic response to closely related hosts was highly host specific, with 90% of the quantifiable transcriptome showing host-dependent expression differences (Figure 3B). This indicates that Botrytis can sense and differentially respond to subtle differences between hosts. Many of the Botrytis genes differentially expressed between hosts encoded cell wall-modifying enzymes (Figure 4A), a core component of Botrytis virulence (van Kan 2006). We identified 27 putative Botrytis CAZymes that were differentially expressed during infection of leaves of closely related hosts (Table S3). In contrast, prior work using Botrytis isolate B05.10 reported that none of the 1,155 CAZymes encoded in the genome showed host-specific expression on lettuce leaves compared with grape and tomato fruit, showing the utility of using an isolate collection (Blanco-Ulate et al. 2014). Our result indicates cell wall-modifying gene expression during leaf infection may exhibit greater host specificity and responsiveness than previously appreciated. In contrast to these host-specific patterns, expression of some Botrytis specialized metabolic pathways were more conserved between hosts. Biosynthetic gene clusters for the phytotoxins botrydial and botcinic acid showed conserved expression across hosts but varied across isolates (Figure 5), indicating induction of these major virulence pathways is similar on these two hosts. Nevertheless, the full breadth of phytotoxic metabolites produced by diverse isolates of Botrytis remains poorly characterized. We also identified several Botrytis specialized metabolic enzymes differentially expressed between hosts (Figure 4A, Figure 4B), and because small changes in pathway composition can substantially alter final metabolite profiles, it is likely that Botrytis deploys distinct blends of specialized metabolites even when infecting closely related hosts. Together, these results indicate that Botrytis balances conserved virulence strategies with finely tuned, host-specific mechanisms to optimize infection across closely related hosts.

The two host species also showed host-dependent differences. Single copy orthologs between common bean and cowpea enabled direct comparison of the host transcriptional responses to Botrytis. Among expressed host single copy orthologs, 97% were differentially expressed between the hosts that was largely quantitative (Figure 10). Despite this extensive divergence, both hosts retained canonical features of the eudicot defense response to necrotrophs. In both hosts, JA and ethylene signaling pathways were upregulated, while photosynthetic machinery was downregulated, consistent with prior studies of Botrytis infection in Arabidopsis (Windram et al. 2012; Zhang et al. 2017; Zhang et al. 2019) and lettuce (De Cremer et al. 2013). Phenylpropanoid metabolism was also induced in both hosts, in line with its well-established role in biotic defense (Ranjan et al. 2019; Ninkuu et al. 2025). Alongside these shared pathway-level responses, many single copy orthologs exhibited opposite regulation between hosts, demonstrating that conserved genes can respond differently to the same pathogen challenge (Figure 8). These oppositely regulated orthologs spanned diverse functions, with no entire pathways showing consistent opposite regulation. Thus, while major host defense pathways are conserved between closely related hosts, this response is tailored with host-specific gene expression.

### Shared pathways swap gene membership according to host variation

Focusing on co-expression networks in both the host and pathogen identified GCNs differentially modulated between the hosts. These had a similar architecture in which a common core set of genes in the GCN were shared across the two hosts, while the extended membership of the module showed extensive gene swapping. This suggests a potential structure for explaining plasticity in host-generalist interactions, whereby instead of full rewiring of pathways depending on the host/attacker, metabolism is shifted around a common core.

Among the Botrytis GCNs, we identified an illustrative example of how gene membership can shift around a common core in a host-dependent manner. Using this example gives an outline of how this particular shift has plausible mechanistic consequences. This Botrytis GCN centers around an NRPS gene cluster on chromosome 12 (Figure 6). NRPS genes produce non-ribosomal peptides, a structurally diverse group of natural products with a broad range of biological activities, including as toxins, pigments, and siderophores (Martínez-Núñez and López 2016). NRPS gene clusters are common in fungi, and previous analysis of the B05.10 genome recovered 11 distinct NRPS clusters (Valero-Jiménez et al. 2020). However, little is described about their potential function in Botrytis virulence. Previously, NRPS1 in B05.10 has been shown to confer protection for Botrytis against exogenous toxins, but its deletion actually increased virulence (Fernández-Morales et al. 2021). The Botrytis GCN when infecting common bean contains the presence of several key flavonoid precursors and both a PKS and NRPS gene cluster. Other fungi have hybrid PKS-NRPS enzymes that catalyze steps in flavonoid biosynthesis (Zhang et al. 2022). In contrast, on cowpea, this Botrytis GCN only consists of the NRPS gene cluster, and all the shikimate/flavonoid pathway genes are absent. This may indicate that this GCN has a host-dependent modularity in which an NRPS gene cluster can be used for divergent purposes by the same pathogen infecting different hosts.

A similar pattern of core GCNs with modified gene membership was also found when comparing the two host species (Figure 9A). One example was a phenylpropanoid associated GCN that shares a common set of genes across the two hosts with a set of distinct genes in each species (Figure 9C). Although the core enzymatic steps of the phenylpropanoid pathway are conserved across land plants, gene expression is extensively regulated by species-specific transcriptional networks in response to environmental stimuli (Liu et al. 2015). Recent support for the ability of plant regulatory networks to rapidly evolve was evidence that angiosperm gene expression patterns evolve more rapidly than animals (Schuster et al. 2026).

The identification of this pattern in both host and pathogen suggests shifting regulatory network membership may play a role in the changing relationship between closely related hosts. Complicating this model are some limitations on the utility of network approaches to fully describe biological systems. In particular, it is critical to emphasize GCNs represent correlations only, and do not measure the distance between interacting genes. Additionally, complexities of gene orthology/paralogy complicates mapping GCNs across distant species. The use of closer related species with a high fraction of singly copy orthologs limits but does not obliviate this complication. While the specific mechanistic hypotheses generated in this study will require further validation, these findings show host-dependent transcriptional responses involve network modulation around a common core of genes in both host and pathogen.

### Disease similarity hides molecular variation in both host and pathogen

Frequently, phenotypic surveys are used prior to conducting molecular assays to identify comparisons with the largest changes. This makes an assumption that virulence phenotype and the molecular underpinnings are largely comparable. The Botrytis-bean and Botrytis-cowpea comparison did not support this assumption. While Botrytis lesion formation was similar in these closely related hosts, there were distinct, complex, hidden gene expression differences in both host and pathogen underlying this phenotype. Therefore, fairly dissimilar transcriptomes can lead to similar disease outcomes. The first line of evidence supporting this is that lesion formation was primarily driven by Botrytis isolate variation, with no significant host x isolate interaction (Figure 2C; Table S1), while a majority of both host and pathogen transcripts had a significant host x isolate interaction (Figure 3B; Figure 10). The second line of evidence is that variation and means for lesion sizes were similar between hosts (Figure 2A), while host and pathogen transcriptomes showed comparably large divergence between hosts. Botrytis transcriptomes were overall highly distinguishable by the host being infected (Figure 3A). Finally, the expression of single copy orthologs between the hosts differed markedly (Figure 10A).

In the context of host generalism, these results caution against assumptions that similar disease outcomes implicitly arise from conserved molecular interactions. Instead, Botrytis achieves phenotypic consistency through flexible, host-specific transcriptional programs, and likewise hosts execute distinct responses that can converge on similar measures of susceptibility. This pattern is consistent with a convergent evolutionary framework, in which similar disease phenotypes can be maintained by selection on pathogenic success, even when the underlying host-pathogen molecular interactions differ substantially. Under this model, host generalism is enabled not by mechanistic uniformity, but by genome-wide transcriptional plasticity in both organisms. We highlight a broader need for co-transcriptome studies to strengthen this model and continue enhancing our understanding of the biology underlying generalist pathogens.

## Materials and Methods

### Plant materials and growth conditions

To test how genetic diversity within closely related hosts of Botrytis influence disease outcome, four genotypes for each of two related host species, common bean (*Phaseolus vulgaris*) and cowpea (*Vigna unguiculata*) were included. Genotypes were sourced from the Gepts lab at UC Davis and included UC Cran, UC Tiger’s Eye, Black Nightfall, and Orca for common bean, and CB27, CB46, Sanzi, and IT97K-499-35 for cowpea. All plants in this study were grown in the same greenhouse with 16 hours of supplemental lighting, 30-80% humidity, and temperature at 65-80℉. Seeds were germinated and grown in Agronomy Mix and drip irrigated 5 times a day. For lesion phenotyping, plants were grown for two independent trials, one in May 2022 and one in July 2022. For transcriptomics, common bean was grown in January 2024 while cowpea was grown in April 2024. Plants were grown for the formation of several true leaves and before flowering, approximately 4 weeks.

### *Botrytis cinerea* materials and growth conditions

A set of 72 Botrytis isolates were used in this study, as described previously (Singh et al. 2025). This set of 72 samples the genetic diversity and virulence of a larger population of 96 Botrytis isolates, originally isolated as single spores from plant tissues collected mainly from California and internationally (Zhang et al. 2017; Zhang et al. 2019; Caseys et al. 2021). There is no evidence of population structure by host or geography in this collection of isolates (Atwell et al, 2018). Isolates were stored at -80°C as a spore stock in 60% glycerol. For experiments, spores were diluted 1:10 (v/v) glycerol stock in sterile grape juice and grown on potato dextrose agar for two weeks to obtain spores for infection assays.

### Detached leaf assay

To measure the virulence of Botrytis and hosts within the legumes, the above eight host genotypes of common bean and cowpea were infected with 72 Botrytis isolates using a previously described detached leaf assay (Denby et al. 2004; Corwin, Copeland, et al. 2016; Soltis et al. 2019). This used a randomized complete block design with 3 independent replicates for each host-genotype combination. Performed across two independent experiments, this yielded 6 replicates of lesion phenotype data for each host-pathogen genotype combination.

For each above independent experiment, leaves were collected from approximately 40 plants for each genotype and randomly distributed across trays containing 1cm of 1% phytoagar with humidity domes at room temperature. Inoculum was generated by growing spores on PDA plates for 10-14 days, collecting spores into water, then diluting them into 50% grape juice to 10 spores/uL. Grape juice was used as the inoculum medium to promote consistent germination across isolates. This alleviates issues where Botrytis has genetic variation impacting germination on single sugar sources (Blakeman 1975; Clark 1977; Benito et al. 1998; Denby et al. 2004). Detached leaves were inoculated on two sites on either side of the adaxial leaf surface (Figure 1) with two different Botrytis isolates.

### Lesion measurement and analysis

To measure the developing lesions, images of trays containing inoculated leaves were collected at 24, 48, 72, and 96 hours after inoculation (HAI). At 72 HAI, lesion formation was visible and quantifiable on leaves. For 72 HAI and 96 HAI images, lesion areas were traced digitally using the EBImage and CRImage packages (Pau et al. 2010; Failmezger et al. 2012) in a previously described R pipeline (Fordyce et al. 2018), then manually inspected to verify accuracy. Lesion areas in pixels were converted to mm^2^ using a reference scale included within each image.

Lesion area was modeled using the lme4 package (Bates et al. 2015). We modeled lesion area according to the host, pathogen, and their interactions while accounting for experimental design with the following:

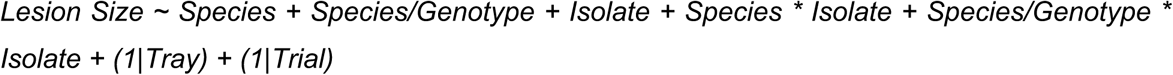

Host genetic diversity, including both “Species” and “Species/Genotype” (genotype nested within species), pathogen genetic diversity as “Isolate”, and their interactions were treated as fixed effects. Experimental design components, including “Tray” (phytoagar flat containing inoculated leaves within a randomized block design) and “Trial” (independent experiment) were treated as random effects.

### RNAseq library preparation and sequencing

To measure the associated gene expression patterns of both host and pathogen during infection, the above detached leaf assay was conducted with 1 selected genotype each from common bean and cowpea. The host genotypes were selected based on a number of factors, including range of infection phenotypes across the Botrytis isolates, available reference genomes, and ease of healthy plant growth. The detached leaf assay was performed as 1 independent experiment for each host species, yielding 3 replicates of transcriptomic data.

At 48 HAI, a 1.20 cm diameter leaf disc containing the inoculation site was removed and immediately flash frozen in liquid nitrogen. Mock leaf samples that were inoculated with grape juice only were also collected at 48 HAI. The 48 HAI timepoint was selected based on a pilot experiment conducted at 24, 30, 36, and 48 HAI, which showed that 48 HAI yielded the highest proportion of fungal-mapped reads. Inoculated leaf discs were stored at -80°C until processing. Libraries were prepared according to a previously described method (Kumar et al. 2012; Zhang et al. 2017). RNA extraction was conducted by placing samples in liquid nitrogen and then homogenizing by rapid agitation in a bead beater followed by direct mRNA isolation using the Dynabeads oligo-dT kit. First and second strand cDNA was produced from the mRNA using an Invitrogen Superscript III kit. The resulting cDNA was fragmented, end-repaired, A-tailed and barcoded as previously described. Adapter-ligated fragments were enriched by PCR and size-selected using AMPure XP beads for a mean of 300 bp prior to sequencing. Barcoded libraries were pooled in batches of 74 (1 host species inoculated with 72 isolates and 2 mocks) and submitted for paired end 150 bp sequencing on a single lane per pool using the AVITI sequencing platform at the UC Davis Genome Center (DNA Technologies Core, Davis, CA).

### Transcriptomic data analysis

The resulting fastq files were separated by adapter index into individual sample files. Quality control was done with MultiQC (Ewels et al. 2016). Raw paired-end sequencing reads were quality-trimmed and filtered using Trimmomatic (Bolger et al. 2014). Adapter sequences were clipped using the ILLUMINACLIP module, allowing up to two seed mismatches, a palindrome clip threshold of 30, and a simple clip threshold of 10. Reads were further processed to remove low-quality bases from the 5’ and 3’ ends, trim within a sliding window of four bases where the average quality dropped below 15, and discard reads shorter than 20 bases. Trimmed and paired reads were retained for downstream analysis, and quality was reassessed with MultiQC. Reads were mapped to reference genomes using Hisat2 (Kim et al. 2019). Bulk reads were mapped first to the host genome (Lonardi et al., 2019; Schmutz et al., 2014). The remaining un-mapped reads were mapped to Botrytis B05.10 isolate reference genome (Van Kan et al. 2017). Read counts were obtained from the mapped reads using RSubread (Shi 2017). Read counts were TMM normalized with the EdgeR package (Robinson et al. 2010). Low expression genes (genes with <1 CPM in <20% of samples within each host) were removed from the dataset. All statistical analyses were conducted within R.

To estimate the effects of host and pathogen on all transcripts, a generalized linear model with a negative binomial distribution (log link) was fitted using the glmmTMB R package. Host gene expression was modeled for each host gene across both species with the following:

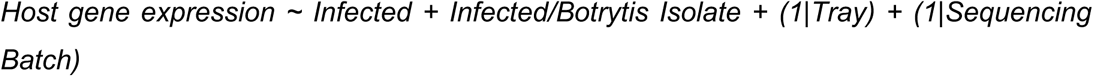

Infection status, including both “Infected” (inoculated with any Botrytis vs mock) and “Infected/Botrytis Isolate” (Botrytis isolate nested within infected) were treated as fixed effects. Experimental design components, including “Tray” (phytoagar flat containing inoculated leaves within a randomized block design) and “Sequencing Batch” (sample set prepared and sequenced together) were treated as random effects.

For the Botrytis transcripts, both hosts were included in the same model. The effects of host species, Botrytis isolate, and their interaction was modeled using the following formula:

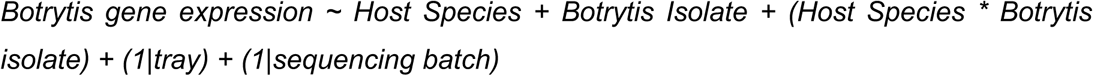

Estimated marginal means and standard error for each transcript were determined for each host genotype & Botrytis isolate using the emmeans package (Lenth and Piaskowski 2017). Model results were summarized using Type II Wald Chi-square tests from the car package to evaluate the significance of each fixed effect. Resulting p-values were adjusted for multiple comparisons using the false discovery rate (FDR) correction (Benjamini and Yekutieli 2001).

Principal component analysis (PCA) was conducted in R’s base package (R Core Team 2024). To evaluate correlation of Botrytis lesion size and Botrytis transcriptome abundance to principal components, the following model was used:

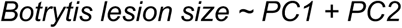

To assess enrichment of certain gene classes in differentially regulated Botrytis genes, hypergeometric tests were performed with the stats package in R. P values were calculated for both overenrichment and underenrichment of the gene class for a given regulation group, then false discovery rate (FDR) adjusted.

### Co-expression network analysis

To assess gene co-expression patterns both within and across host and pathogen, we used a previously described gene co-expression network analysis pipeline (Wisecaver et al. 2017). For each host–pathogen combination, normalized gene counts from expressed genes were estimated separately from host and pathogen transcripts. Normalized host and pathogen transcriptomes were then combined and used to calculate correlations among all transcripts. The pipeline was run twice: once for common bean infected with Botrytis and once for cowpea infected with Botrytis. Pearson correlation coefficient (PCC) values for each gene pair were calculated, then scored for mutual rank. Mutual ranks were used to call modules of tightly coexpressed genes using ClusterONE with a decay rate of 5 (Nepusz et al. 2012). Networks with at least 10 gene members and a significant network correlation statistic (p < 0.05) were retained for further analysis. This yielded 378 total host & Botrytis networks on common bean, and 428 total networks of host & Botrytis networks on cowpea. Across this set of networks, the average number of genes per network was 18. For gene co-expression networks of interest, overall network expression level was calculated by z-scaling expression values within each gene in the dataset and taking the mean expression across genes in the network.

### Host legume orthology analysis

To compare responses to Botrytis directly between the two host species, orthology analysis was conducted with Orthofinder (Emms and Kelly 2019). Briefly, this analysis used predicted proteomes from each host species to identify orthogroups across the two species. The algorithm then infers gene trees, a rooted species tree, then analyzes the rooted gene trees to identify orthologs and gene duplication events. This results in a full list of predicted orthologs between the species, including single copy orthologs and any duplicated genes.

To estimate the main functions of host single copy orthologs from Orthofinder grouped by their response to infection vs. mock, gene ontology (GO) enrichment analysis was performed using the R package topGO (Adrian Alexa 2017). This involved four separate tests: GO enrichment of genes upregulated in both hosts, genes downregulated in both hosts, genes upregulated in common bean but downregulated in cowpea, and genes upregulated in cowpea but downregulated in common bean. The analysis tested for enrichment of terms from the Biological Process ontology among differentially expressed single copy orthologs, using all single copy orthologs as the background. Enrichment was assessed with Fisher’s exact test under the classic algorithm, and terms were ranked by p-values.

To assess the relative contribution of host and Botrytis genetic diversity to the expression of host single copy orthologs, a generalized linear model with a negative binomial distribution (log link) was fitted using the glmmTMB R package:

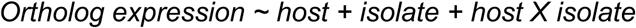

Model significance was evaluated using Type II Wald Chi-square tests implemented in the car package. Resulting p-values were adjusted for multiple comparisons using the FDR correction (Benjamini and Yekutieli 2001). Variance components were calculated from sums of squares.

## Acknowledgements

This work was supported by the NSF award IOS 2020754 for DJK. We thank the Gepts and Diepenbrock labs at UC Davis for providing seeds, all members of the Kliebenstein lab for their support, and HPC@UCD for providing computational resources that have contributed to the research results reported in this paper.

## Author contributions

D.J.K.: designed research; A.J.M., R.S., C.T., K.S., L.F., B.G., and C.C.: performed research; A.J.M.: analyzed data; A.J.M.: Writing—original draft,; D.J.K., R.S.: Writing—review and editing.

## Data availability

All data supporting this study are included in the main article and/or the supplementary information. The RNA-seq data are uploaded to the NCBI Sequence Read Archive (SRA) under BioProject ID [PRJNA1217477], and will be made publicly available upon publication.

## Supplementary Figures

**Figure S1.**
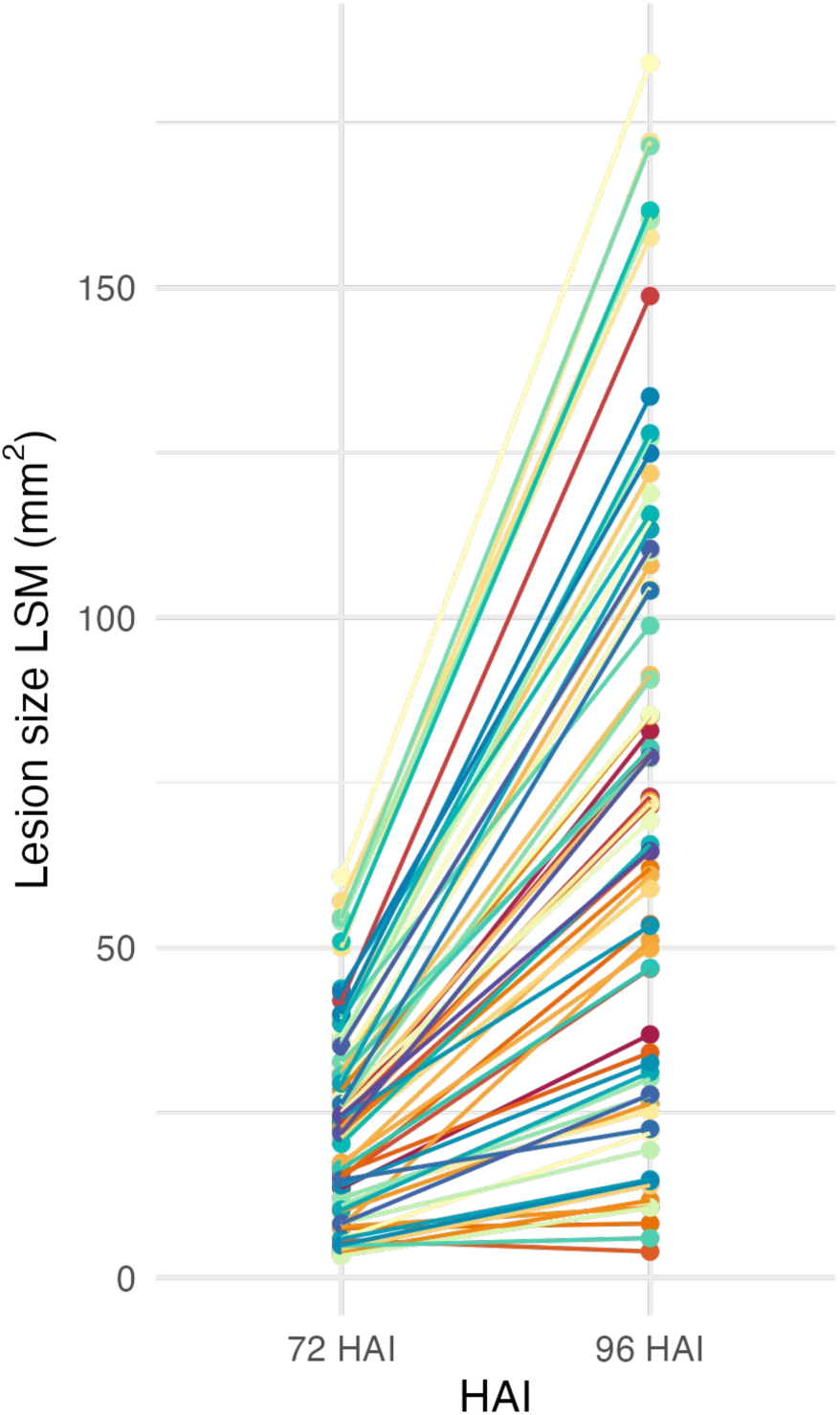
Least-squared means of lesion sizes of 72 Botrytis isolates on legume hosts over time. Each isolate is shown as a different colored line and its lesion value is averaged across 8 host genotypes. HAI = Hours after inoculation.

**Figure S2.**
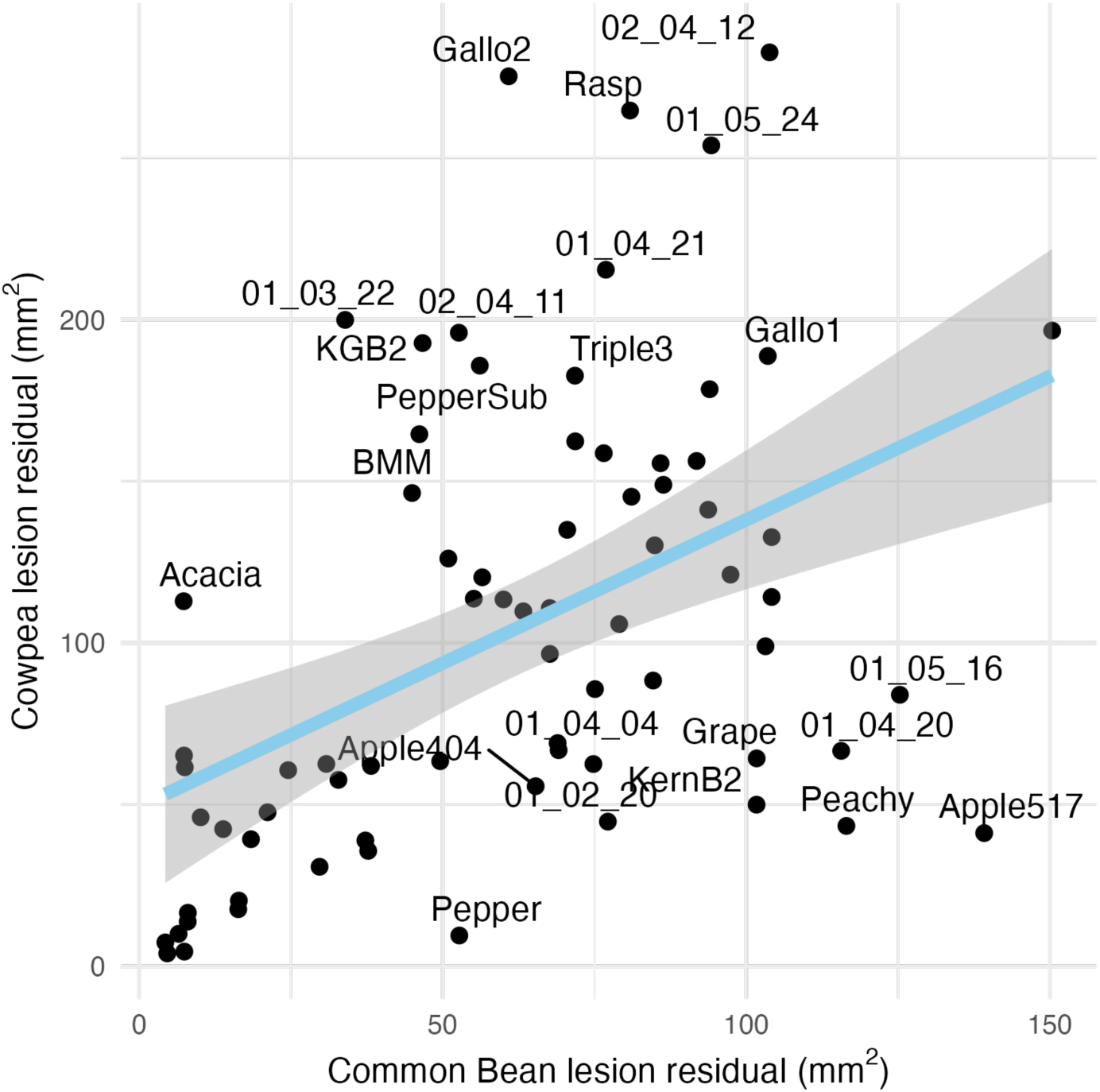
Comparative lesion size residuals of specific isolates as a metric for specialism of each isolate. Genotype residuals were calculated as the mean of one isolate on a given genotype subtracted from the overall mean of that isolate across the species. Species-level residuals were then calculated for each isolate by taking the absolute value sum of the four genotype residuals for each species. Outlying isolates are labeled with the isolate name. Correlation statistics: R^2^ = 0.458, p < 0.01.

**Figure S3.**
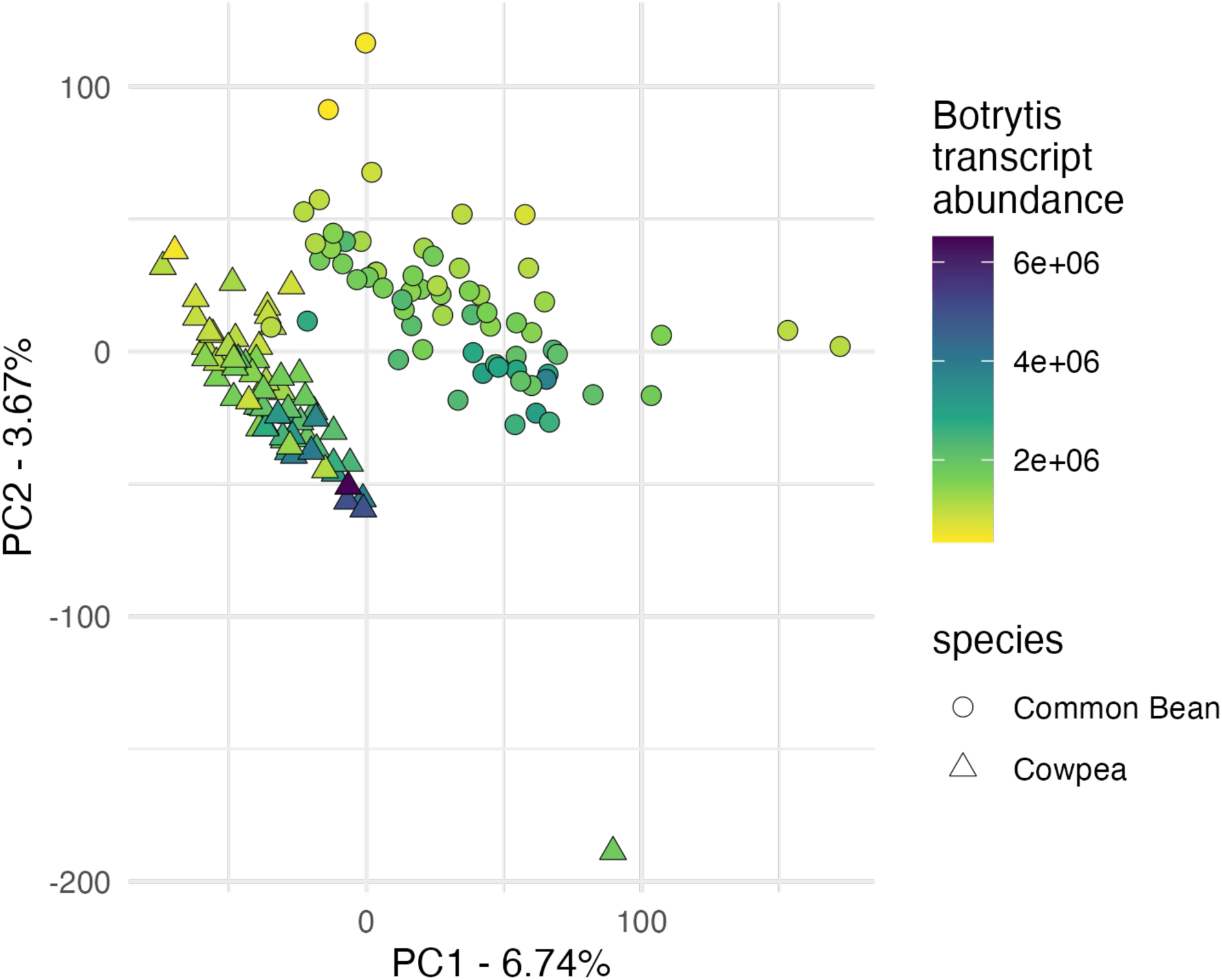
PCA of overall Botrytis transcriptome at 48 HAI. Shape of points show the species the infecting Botrytis isolate was collected from, where circles are common bean and triangles are cowpea. Points are colored by total transcript abundance of that isolate on that host species at 48 HAI.

**Figure S4.**
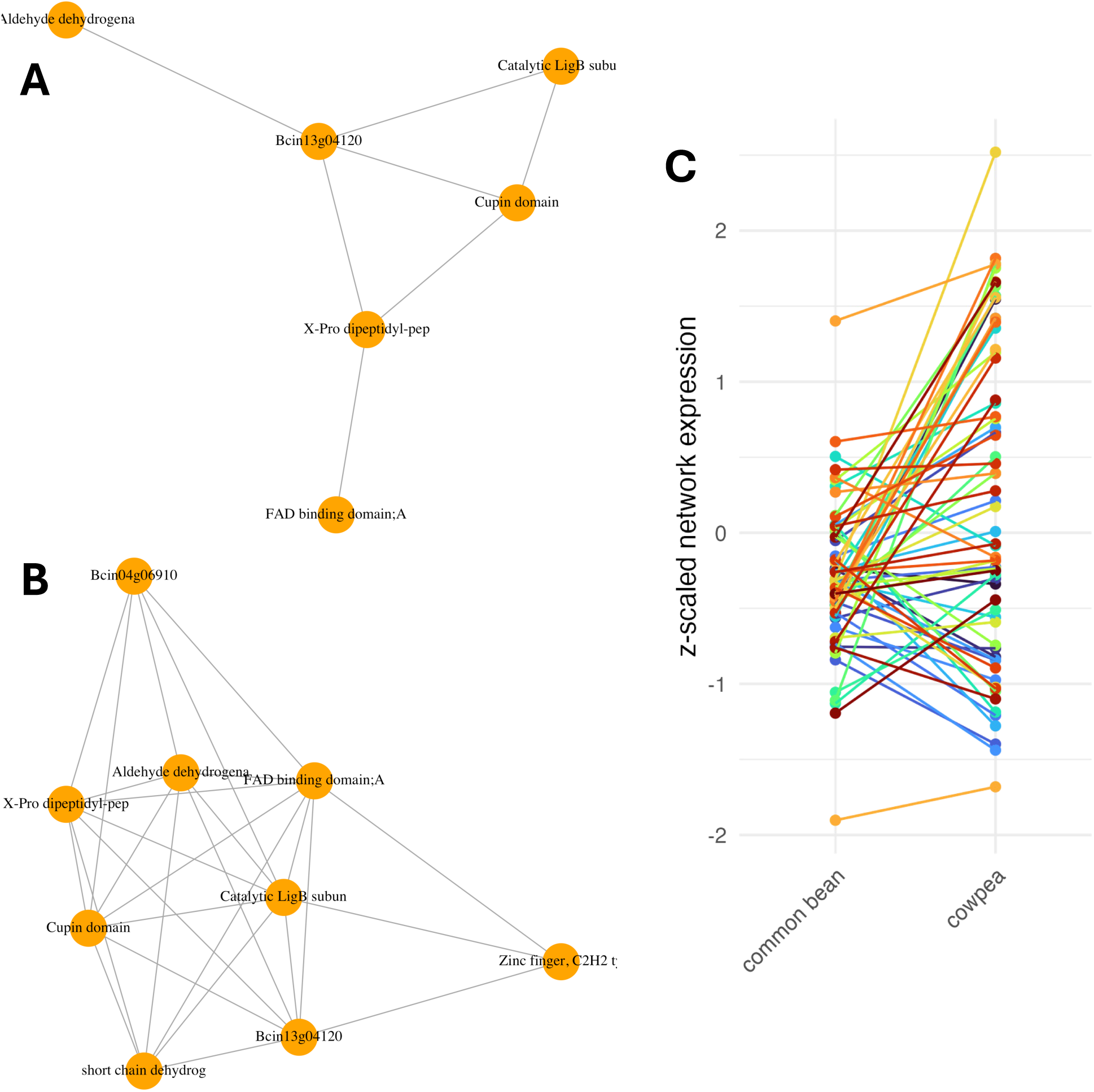
Selected Botrytis co-expression networks containing several Botrytis genes with significant host x isolate interaction. Similar Botrytis gene co-expression networks were detected when infecting both A) common bean and B) cowpea with slight differences. The network represents co-expression of a Botrytis gene cluster on chromosome 13. C) Z-scaled expression of these interaction networks on different hosts. Colored points and connecting lines represent different Botrytis isolates.

**Figure S5.**
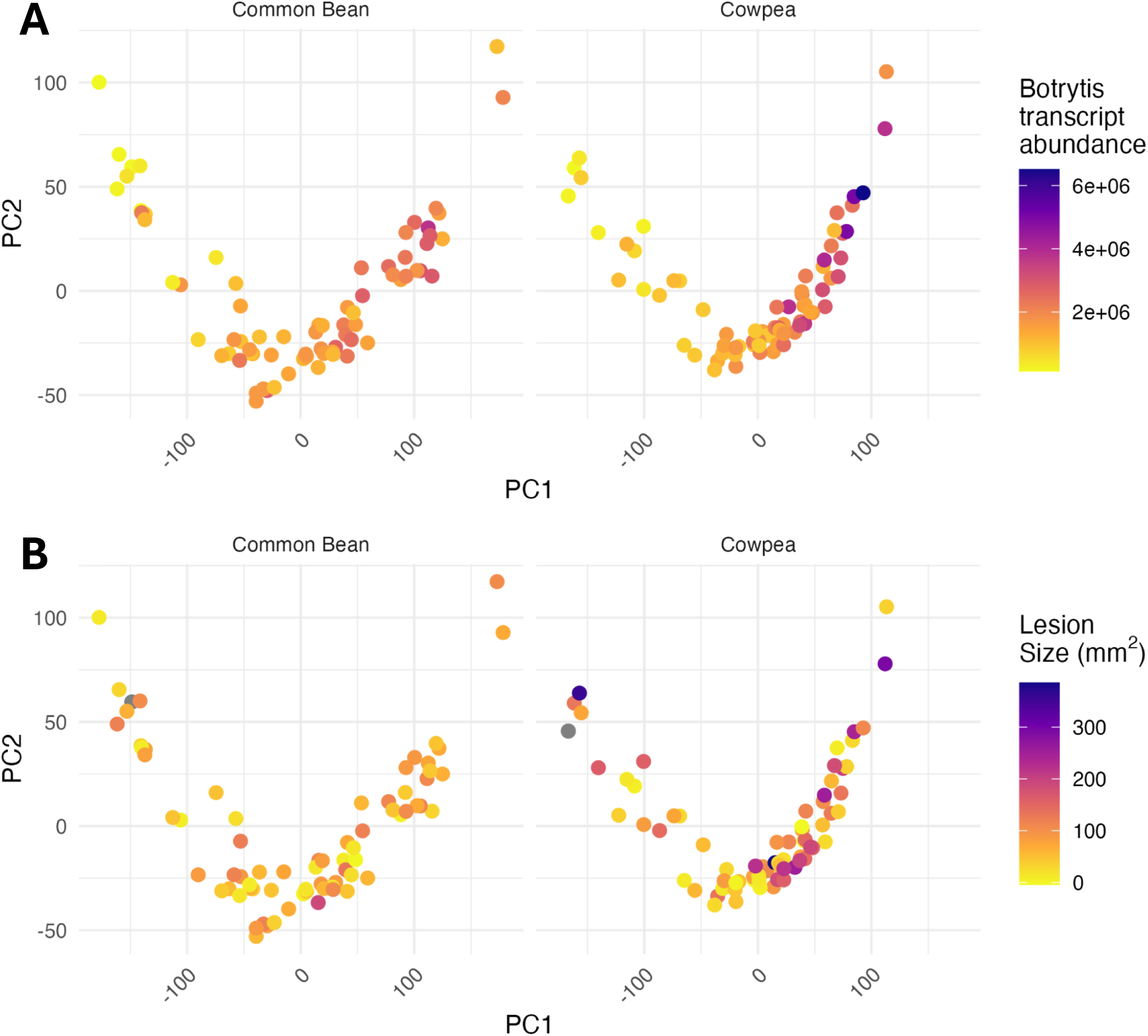
PCA of host transcriptomes colored by A) Botrytis transcript abundance and B) lesion size at 96 HAI in mm^2^. Each point represents a Botrytis isolate on either common bean or cowpea. Isolate colored grey represents the mock inoculation. PC1 accounts for 42% of the variance in cowpea, 39% variance in common bean; PC2 accounts for 8% of the variance in cowpea, 7% variance in common bean.

**Figure S6.**
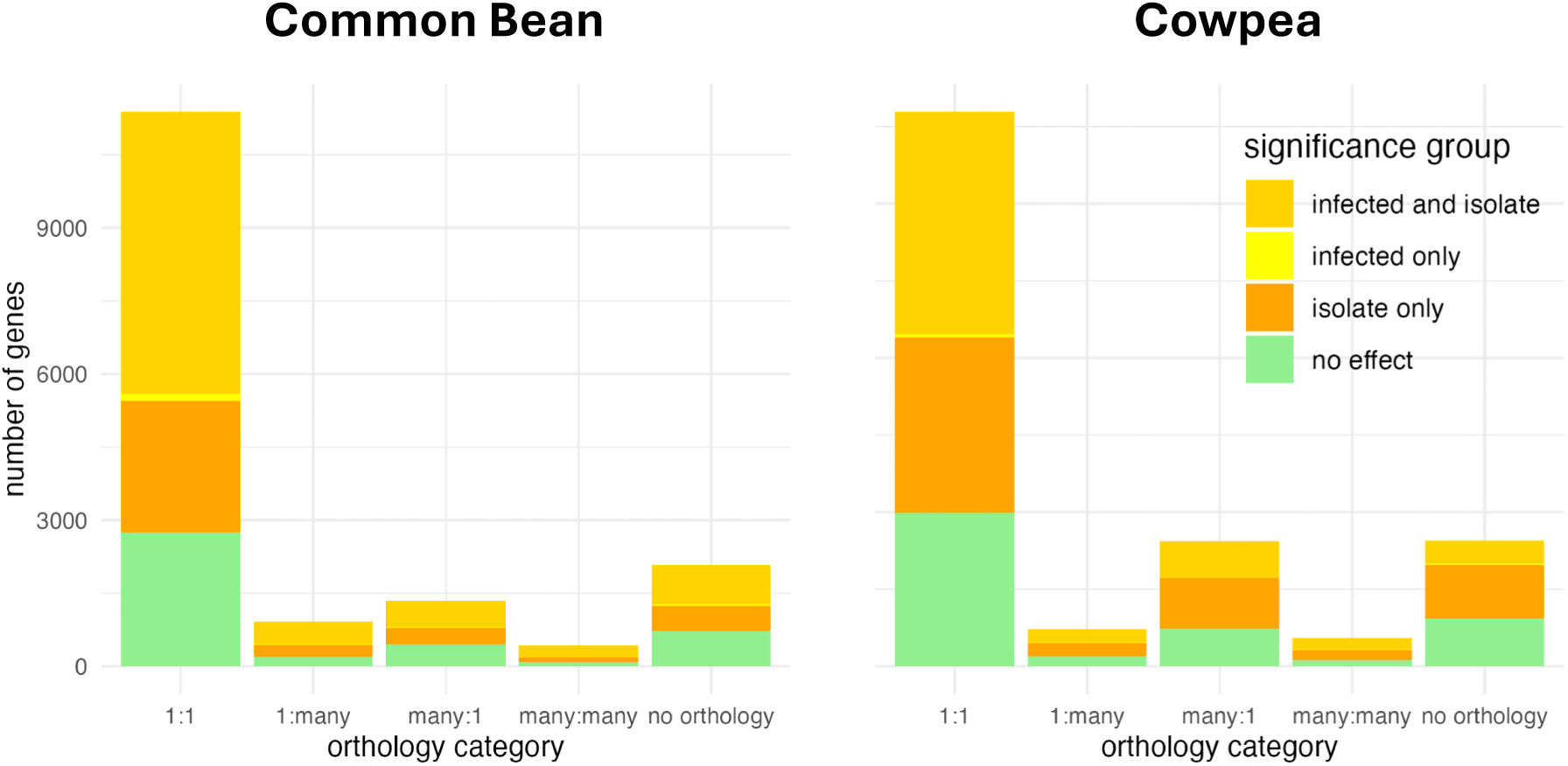
Expressed gene ortholog summary between common bean and cowpea. Single copy (1:1) orthologs are those that have a single match in either legume species, while 1:many orthologs are those that have one gene in one species that match >1 in the other species, etc. Gene counts are colored by their significance in the model gene expression ∼ infected + infected/isolate. “Infected only” are genes only significant in the *infected* main effect. “Isolate only” are genes only significant in the *infected/isolate* (nested) term. “Infected and isolate” are genes that are significant for both the *infected* main effect and the *infected/isolate* term.

## Supporting Information

Additional supporting information may be found in the online version of this article.

**Table S1** ANOVA of lesion sizes during *Botrytis cinerea* infection of *Phaseolus vulgaris* and *Vigna unguiculata*.

**Table S2** GO enrichment of the 58 solely host effect genes against the whole *Botrytis cinerea* genome when infecting *Phaseolus vulgaris* and *Vigna unguiculata*.

**Table S3** *Botrytis cinerea* genes differentially expressed across 2 host legume species (*Phaseolus vulgaris* and *Vigna unguiculata*).

**Table S4** 18 identified *Botrytis cinerea* GCNs with isolate specificity, but similarly expressed across hosts (*Phaseolus vulgaris* and *Vigna unguiculata*).

**Table S5** Gene lists for the main phytoxic gene clusters in *Botrytis cinerea*, botrydial and botcinic acid.

**Table S6** Lesion sizes and mean expression level of key *Botrytis cinerea* phytotoxin clusters for each combination of *B. cinerea* isolate and host species (*Phaseolus vulgaris* and *Vigna unguiculata*).

**Table S7** GO enrichment of legume host single copy orthologs in *Phaseolus vulgaris* and *Vigna unguiculata* when infected with *Botrytis cinerea*.

**Table S8** 37 *Botrytis cinerea* infection-responsive host GCNs containing >=50% single copy orthologs between common bean (*Phaseolus vulgaris*) and cowpea (*Vigna unguiculata*).

## References

1. Adrian Alexa JR. 2017. topGO. [accessed 2025 Nov 10]. https://bioconductor.org/packages/topGO.

2. Baetsen-Young A, Man Wai C, VanBuren R, Day B. 2020. *Fusarium virguliform e* Transcriptional Plasticity Is Revealed by Host Colonization of Maize versus Soybean. Plant Cell. 32(2):336–351.

3. Barrett LG, Heil M. 2012. Unifying concepts and mechanisms in the specificity of plant–enemy interactions. Trends Plant Sci. 17(5):282–292.

4. Bates D, Mächler M, Bolker B, Walker S. 2015. Fitting Linear Mixed-Effects Models Using **lme4**. J Stat Softw. 67(1) [accessed 2025 Nov 5].

5. Benito EP, Ten Have A, Van ’T Klooster JW, Van Kan JAL. 1998. Fungal and plant gene expression during synchronized infection of tomato leaves by Botrytis cinerea. Eur J Plant Pathol. 104(2):207–220.

6. Benjamini Y, Yekutieli D. 2001. The control of the false discovery rate in multiple testing under dependency. Ann Stat. 29(4) [accessed 2025 Aug 17].

7. Blakeman JP. 1975. Germination of Botrytis cinerea conidia in vitro in relation to nutrient conditions on leaf surfaces. Trans Br Mycol Soc. 65(2):239–247.

8. Blanco-Ulate B et al. 2014. Genome-wide transcriptional profiling of Botrytis cinerea genes targeting plant cell walls during infections of different hosts. Front Plant Sci. 5 [accessed 2025 Nov 11].

9. Bolger AM, Lohse M, Usadel B. 2014. Trimmomatic: a flexible trimmer for Illumina sequence data. Bioinformatics. 30(15):2114–2120.

10. Caseys C et al. 2021. Quantitative interactions: the disease outcome of *Botrytis cinerea* across the plant kingdom. G3 GenesGenomesGenetics. 11(8):jkab175.

11. Clark CA. 1977. Comparative Nutrient Dependency of Botrytis squamosa and B. cinerea for Germination of Conidia and Pathogenicity on Onion Leaves. Phytopathology. 77(2):212.

12. Colmenares AJ et al. 2002. The Putative Role of Botrydial and Related Metabolites in the Infection Mechanism of Botrytis cinerea. J Chem Ecol. 28(5):997–1005.

13. Corwin JA, Copeland D, et al. 2016. The Quantitative Basis of the Arabidopsis Innate Immune System to Endemic Pathogens Depends on Pathogen Genetics Koenig D, editor. PLOS Genet. 12(2):e1005789.

14. Corwin JA, Subedy A, Eshbaugh R, Kliebenstein DJ. 2016. Expansive Phenotypic Landscape of *Botrytis cinerea* Shows Differential Contribution of Genetic Diversity and Plasticity. Mol Plant-Microbe Interactions®. 29(4):287–298.

15. Dalmais B et al. 2011. The Botrytis cinerea phytotoxin botcinic acid requires two polyketide synthases for production and has a redundant role in virulence with botrydial: Botcinic acid biosynthesis gene clusters. Mol Plant Pathol. 12(6):564–579.

16. De Cremer K et al. 2013. RNA seq-based transcriptome analysis of *L actuca sativa* infected by the fungal necrotroph *Botrytis cinerea* . Plant Cell Environ. 36(11):1992–2007.

17. Denby KJ, Kumar P, Kliebenstein DJ. 2004. Identification of *Botrytis cinerea* susceptibility loci in *Arabidopsis thaliana*. Plant J. 38(3):473–486.

18. Elad Y, Pertot I, Cotes Prado AM, Stewart A. 2016. Plant Hosts of Botrytis spp. In: Fillinger S, Elad Y, editors. Botrytis – the Fungus, the Pathogen and its Management in Agricultural Systems. Springer International Publishing; p 413–486 [accessed 2025 Nov 19].

19. Emms DM, Kelly S. 2019. OrthoFinder: phylogenetic orthology inference for comparative genomics. Genome Biol. 20(1):238.

20. Ewels P, Magnusson M, Lundin S, Käller M. 2016. MultiQC: summarize analysis results for multiple tools and samples in a single report. Bioinformatics. 32(19):3047–3048.

21. Failmezger H, Yuan Y, Rueda O, Markowetz F. 2012. CRImage: CRImage a Package to Classify Cells and Calculate Tumour Cellularity.

22. Fekete É et al. 2012. Genetic diversity of a Botrytis cinerea cryptic species complex in Hungary. Microbiol Res. 167(5):283–291.

23. Fernández-Morales A et al. 2021. Deletion of the Bcnrps1 Gene Increases the Pathogenicity of Botrytis cinerea and Reduces Its Tolerance to the Exogenous Toxic Substances Spermidine and Pyrimethanil. J Fungi. 7(9):721.

24. Fordyce RF et al. 2018. Digital Imaging Combined with Genome-Wide Association Mapping Links Loci to Plant-Pathogen Interaction Traits. Plant Physiol. 178(3):1406–1422.

25. van Kan JAL. 2006. Licensed to kill: the lifestyle of a necrotrophic plant pathogen. Trends Plant Sci. 11(5):247–253.

26. Kim D et al. 2019. Graph-based genome alignment and genotyping with HISAT2 and HISAT-genotype. Nat Biotechnol. 37(8):907–915.

27. Krishnan P et al. 2023. Polygenic pathogen networks influence transcriptional plasticity in the Arabidopsis–Botrytis pathosystem Birchler J, editor. GENETICS. 224(3):iyad099.

28. Kumar R et al. 2012. A High-Throughput Method for Illumina RNA-Seq Library Preparation. Front Plant Sci. 3 [accessed 2024 Dec 12].

29. Kusch S et al. 2022. Transcriptional response to host chemical cues underpins the expansion of host range in a fungal plant pathogen lineage. ISME J. 16(1):138–148.

30. Leisen T et al. 2022. Multiple knockout mutants reveal a high redundancy of phytotoxic compounds contributing to necrotrophic pathogenesis of Botrytis cinerea Wang Y, editor. PLOS Pathog. 18(3):e1010367.

31. Lenth RV, Piaskowski J. 2017. emmeans: Estimated Marginal Means, aka Least-Squares Means. 2.0.0 [accessed 2025 Nov 5].

32. Li X, Baudry J, Berenbaum MR, Schuler MA. 2004. Structural and functional divergence of insect CYP6B proteins: From specialist to generalist cytochrome P450. Proc Natl Acad Sci. 101(9):2939–2944.

33. Liu J, Osbourn A, Ma P. 2015. MYB Transcription Factors as Regulators of Phenylpropanoid Metabolism in Plants. Mol Plant. 8(5):689–708.

34. Liu Z et al. 2021. The Roles of Cruciferae Glucosinolates in Disease and Pest Resistance. Plants. 10(6):1097.

35. López-Cruz J et al. 2017. Absence of Cu–Zn superoxide dismutase BCSOD1 reduces *Botrytis cinerea* virulence in Arabidopsis and tomato plants, revealing interplay among reactive oxygen species, callose and signalling pathways. Mol Plant Pathol. 18(1):16–31.

36. Martínez-Núñez MA, López VELY. 2016. Nonribosomal peptides synthetases and their applications in industry. Sustain Chem Process. 4(1):13.

37. Mercier A et al. 2021. Population Genomics Reveals Molecular Determinants of Specialization to Tomato in the Polyphagous Fungal Pathogen *Botrytis cinerea* in France. Phytopathology®. 111(12):2355–2366.

38. Moghaddam SM et al. 2021. The tepary bean genome provides insight into evolution and domestication under heat stress. Nat Commun. 12(1):2638.

39. Monte I. 2023. Jasmonates and salicylic acid: Evolution of defense hormones in land plants. Curr Opin Plant Biol. 76:102470.

40. Nepusz T, Yu H, Paccanaro A. 2012. Detecting overlapping protein complexes in protein-protein interaction networks. Nat Methods. 9(5):471–472.

41. Ninkuu V et al. 2022. Lignin and Its Pathway-Associated Phytoalexins Modulate Plant Defense against Fungi. J Fungi. 9(1):52.

42. Ninkuu V et al. 2025. Phenylpropanoids metabolism: recent insight into stress tolerance and plant development cues. Front Plant Sci. 16:1571825.

43. Pau G et al. 2010. EBImage—an R package for image processing with applications to cellular phenotypes. Bioinformatics. 26(7):979–981.

44. Perez De Souza L et al. 2019. Multi-tissue integration of transcriptomic and specialized metabolite profiling provides tools for assessing the common bean (*Phaseolus vulgaris*) metabolome. Plant J. 97(6):1132–1153.

45. R Core Team. 2024. R: A language and environment for statistical computing. https://www.R-project.org/

46. Ranjan A et al. 2019. Resistance against *Sclerotinia sclerotiorum* in soybean involves a reprogramming of the phenylpropanoid pathway and up-regulation of antifungal activity targeting ergosterol biosynthesis. Plant Biotechnol J. 17(8):1567–1581.

47. Robinson MD, McCarthy DJ, Smyth GK. 2010. edgeR : a Bioconductor package for differential expression analysis of digital gene expression data. Bioinformatics. 26(1):139–140.

48. Rowe HC, Kliebenstein DJ. 2007. Elevated Genetic Variation Within Virulence-Associated *Botrytis cinerea* Polygalacturonase Loci. Mol Plant-Microbe Interactions®. 20(9):1126–1137.

49. Schumacher J et al. 2012. Natural Variation in the VELVET Gene bcvel1 Affects Virulence and Light-Dependent Differentiation in Botrytis cinerea Harris S, editor. PLoS ONE. 7(10):e47840.

50. Schuster C et al. 2026. Evolutionary transcriptomics unveils rapid changes of gene expression patterns in flowering plants. Cell. S009286742501428X.

51. Shi W. 2017. Rsubread. [accessed 2025 Nov 5]. https://bioconductor.org/packages/Rsubread.

52. Singh R et al. 2025. Combined generalist and host-specific transcriptional strategies enable host generalism in the fungal pathogen Botrytis cinerea. [accessed 2025 July 29]. http://biorxiv.org/lookup/doi/10.1101/2025.07.24.666639.

53. Soltis NE et al. 2019. Interactions of Tomato and *Botrytis cinerea* Genetic Diversity: Parsing the Contributions of Host Differentiation, Domestication, and Pathogen Variation. Plant Cell. 31(2):502–519.

54. Stefanato FL et al. 2009. The ABC transporter BcatrB from *Botrytis cinerea* exports camalexin and is a virulence factor on *Arabidopsis thaliana*. Plant J. 58(3):499–510.

55. Suárez I, Collado IG, Garrido C. 2024. Revealing Hidden Genes in Botrytis cinerea: New Insights into Genes Involved in the Biosynthesis of Secondary Metabolites. Int J Mol Sci. 25(11):5900.

56. Sucher J et al. 2020. Phylotranscriptomics of the Pentapetalae Reveals Frequent Regulatory Variation in Plant Local Responses to the Fungal Pathogen *Sclerotinia sclerotiorum*. Plant Cell. 32(6):1820–1844.

57. Valero-Jiménez CA et al. 2020. Dynamics in Secondary Metabolite Gene Clusters in Otherwise Highly Syntenic and Stable Genomes in the Fungal Genus *Botrytis* Li-Jun M, editor. Genome Biol Evol. 12(12):2491–2507.

58. Van Kan JAL et al. 2017. A gapless genome sequence of the fungus *Botrytis cinerea*. Mol Plant Pathol. 18(1):75–89.

59. Voisin A-S et al. 2014. Legumes for feed, food, biomaterials and bioenergy in Europe: a review. Agron Sustain Dev. 34(2):361–380.

60. Walker A et al. 2015. Population structure and temporal maintenance of the multihost fungal pathogen *Botrytis cinerea* : causes and implications for disease management. Environ Microbiol. 17(4):1261–1274.

61. Wang C, Liu Y, Li S-S, Han G-Z. 2015. Insights into the Origin and Evolution of the Plant Hormone Signaling Machinery. Plant Physiol. 167(3):872–886.

62. Wang Y et al. 2021. Regulation and Function of Defense-Related Callose Deposition in Plants. Int J Mol Sci. 22(5):2393.

63. Windram O et al. 2012. *Arabidopsis* Defense against *Botrytis cinerea* : Chronology and Regulation Deciphered by High-Resolution Temporal Transcriptomic Analysis. Plant Cell. 24(9):3530–3557.

64. Wink M. 2013. Evolution of secondary metabolites in legumes (Fabaceae). South Afr J Bot. 89:164–175.

65. Wisecaver JH et al. 2017. A Global Coexpression Network Approach for Connecting Genes to Specialized Metabolic Pathways in Plants. Plant Cell. 29(5):944–959.

66. Zhang H et al. 2022. A fungal NRPS-PKS enzyme catalyses the formation of the flavonoid naringenin. Nat Commun. 13(1):6361.

67. Zhang W et al. 2017. Plastic Transcriptomes Stabilize Immunity to Pathogen Diversity: The Jasmonic Acid and Salicylic Acid Networks within the Arabidopsis/ *Botrytis* Pathosystem. Plant Cell. 29(11):2727–2752.

68. Zhang W et al. 2019. Plant–necrotroph co-transcriptome networks illuminate a metabolic battlefield. eLife. 8:e44279.

69. Zhu W et al. 2017. BcXYG1, a Secreted Xyloglucanase from *Botrytis cinerea*, Triggers Both Cell Death and Plant Immune Responses. Plant Physiol. 175(1):438–456.

